# PRSS2 stimulates tumor growth by remodeling the TME via repression of Tsp1

**DOI:** 10.1101/2021.03.23.436667

**Authors:** Lufei Sui, Suming Wang, Debolina Ganguly, Tina El Rayes, Cecilie Askeland, Astrid Børretzen, Danielle Sim, Ole Johan Halvorsen, Gøril Knutsvik, Sura Aziz, Svein Haukaas, William D Foulkes, Diane R. Bielenberg, Arturas Ziemys, Vivek Mittal, Rolf A. Brekken, Lars A. Akslen, Randolph S. Watnick

## Abstract

In the earliest stages of tumor development, epithelial tumors (carcinomas) are physically confined to the area of the tissue in which they form. These nascent lesions (carcinomas in situ) are sequestered from the tissue parenchyma by the basement membrane. Within the tissue parenchyma lie a myriad of cell types comprised of fibroblasts, immune and inflammatory cells and endothelial cells. Upon invasion across the basement membrane and into the tissue parenchyma, tumors must manipulate the expression of pro- and anti-tumorigenic proteins such that pro-tumorigenic factors are produced in excess to anti-tumorigenic proteins. One such anti-tumorigenic protein is Thrombospondin-1 (Tsp-1). We have previously demonstrated that stimulation of Tsp-1 in the tumor microenvironment (TME) potently inhibits tumor growth and progression and in some cases induces tumor regression. Here, we identify a novel tumor-mediated mechanism to repress the expression of Tsp-1 in the TME via secretion of the serine protease PRSS2. We demonstrate that PRSS2 represses Tsp-1, not via its enzymatic activity, but by binding to low-density lipoprotein receptor-related protein 1 (LRP1). These findings describe a novel activity for PRSS2 through binding to LRP1 and represent a potential therapeutic strategy to treat cancer by blocking the PRSS2-mediated repression of Tsp-1.

## Introduction

The paracrine, juxtacrine and exocrine signaling between tumor cells and the nontransformed cells that constitute the tumor microenvironment (TME) are among the key drivers of tumor progression and metastasis. These intercellular signaling pathways regulate such crucial processes as tumor cell invasion and migration, angiogenesis, and immune and inflammatory cell infiltration^1–3^. Thus, the ability of a tumor to alter the activity of the cells in the microenvironment is critical for growth at the primary and metastatic sites ^4–8^. Like many intracellular processes, the balance between extracellular tumor-promoting factors and tumor-inhibitory factors ultimately determines whether tumors grow and expand beyond the primary site or remain localized or even dormant. For example, pro-invasive proteases can be counteracted by protease inhibitors. Similarly, pro-angiogenic factors, such as VEGF, are balanced by anti-angiogenic factors, such as endostatin and thrombospondin-1 (Tsp-1).

Interestingly, tumor-secreted molecules often promote, or inhibit, tumor growth via more than one mechanism. For example, VEGF was identified as a pro-angiogenic factor ^9,10^, but has subsequently been demonstrated to also be a potent immunosuppressive factor ^11–14^. Tsp-1, on the other hand, is a potent inhibitor of angiogenesis, but also mediates the resolution of inflammation by promoting M1 polarization of macrophages^15^. We have previously identified a tumor-secreted protein, prosaposin, (PSAP) that functions as a paracrine inhibitor of primary and metastatic tumor growth ^5^. PSAP inhibits tumor growth and metastasis primarily via the stimulation of the expression of Tsp-1 in myeloid derived suppressor cells (MDSCs) ^4,16^.

In the studies that identified prosaposin as a stimulator of Tsp-1, we also observed the ability of highly metastatic tumor cells to repress Tsp-1 in the TME ^5^. Here, we report the use of a proteomic screen to identify the tumor-secreted, metastasis-promoting, repressor of Tsp-1 as the serine protease PRSS2. PRSS2 is alternatively known as Trypsin-2 and Tumor Associated Trypsin and its overexpression has been linked to, and can induce, pancreatitis^17–19^. Further, we demonstrate that PRSS2 represses Tsp-1, not via its protease activity, but by binding to low-density lipoprotein receptor related protein 1 (LRP1). Strikingly, knockdown of PRSS2 in tumor cells and knockout of LRP1 in myeloid cells dramatically inhibit tumor growth in mouse models of breast and pancreatic cancer, with tumors having higher levels of Tsp-1 in the TME. These findings establish a novel intercellular signaling pathway that stimulates tumor growth and progression via the repression of Tsp-1. Moreover, we have demonstrated that stimulation of Tsp-1 via ectopic expression of PSAP and systemic delivery of a therapeutic peptide derived from PSAP potently inhibits primary and metastatic tumor growth in multiple tumor models ^4,5,16^. Significantly, the anti-tumor therapeutic strategy of augmenting Tsp-1 expression by the PSAP peptide has also been validated in clinical trials ^20,21^. Thus, based on the paracrine mechanism of the anti-tumor activity of Tsp-1, a therapeutic strategy that disrupts its repression by PRSS2 should be indication agnostic.

## Results

### Metastatic tumors repress Tsp-1 in the tumor microenvironment

We have previously reported that highly metastatic human breast and prostate tumor cells derived from weakly metastatic cell lines via serial *in vivo* passaging repress the expression of the anti-tumorigenic protein Tsp-1 in the TME ^5^. To determine whether the repression of Tsp-1 is mediated directly by a tumor-secreted protein(s) we harvested conditioned media (CM) from the highly metastatic prostate cancer cell line PC3M-LN4 prostate cancer along with the weakly metastatic parental cell line PC3 and used them to treat primary human lung fibroblasts. We found that only the CM from the metastatic PC3M-LN4 cell line repressed the expression of Tsp-1 in lung fibroblasts (Figure 1A and B).

**Figure 1.**
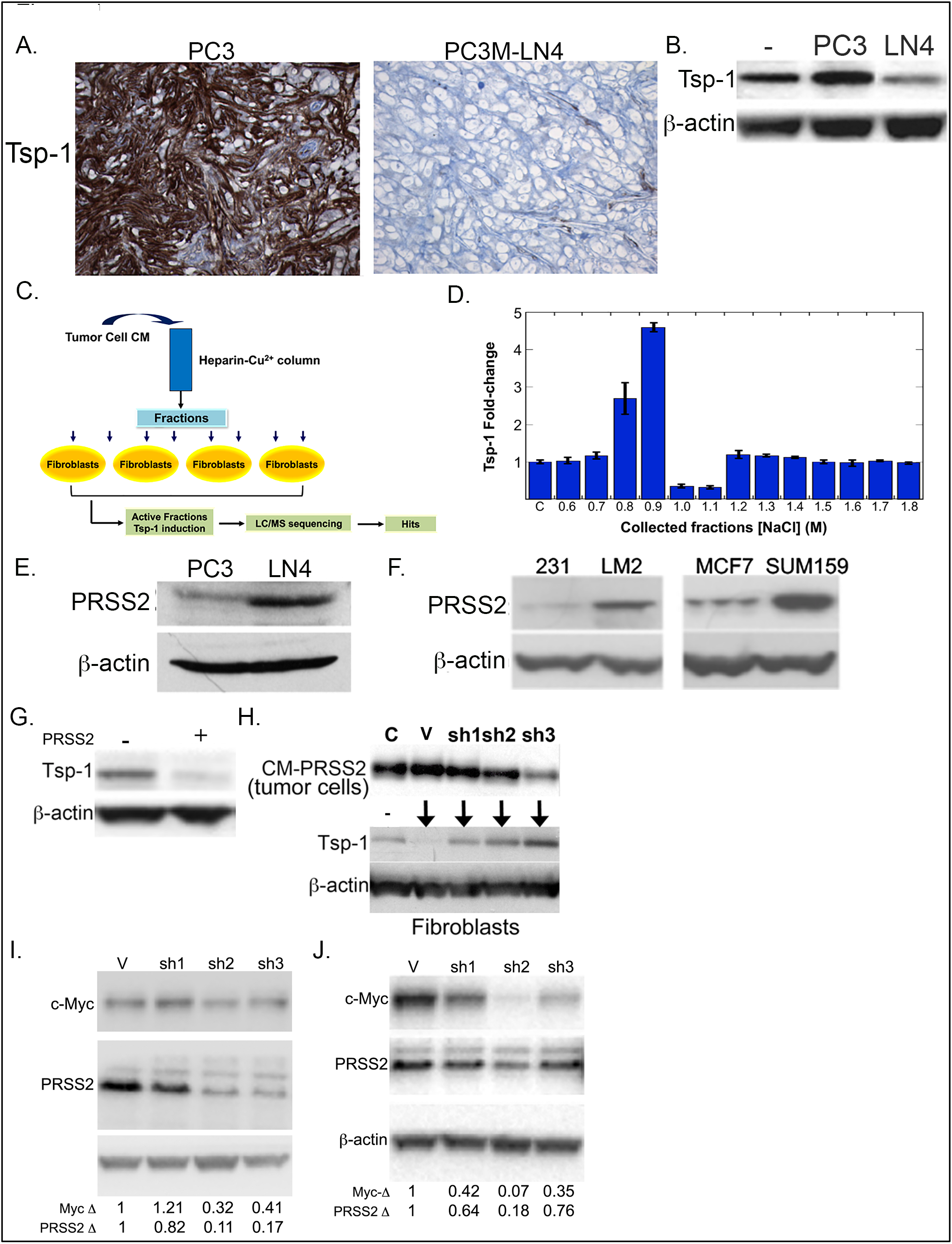
Breast and prostate cancer cells repress Tsp-1 in a paracrine manner via PRSS2. **(A)** Immunohistochemistry (IHC) of thrombospondin-1 (Tsp-1) expression in tumors formed in the prostate gland of SCID mice by PC3 and PC3M-LN4 prostate cancer cells; **(B)** Western blot of Tsp-1 and actin expression in WI-38 fibroblasts that were untreated (-) or treated with CM from PC3 and PC3M-LN4 (LN4) prostate cancer cells; **(C)** Schematic for the identification of Tsp-1 repressing protein in PC3M-LN4 conditioned media; **(D)** ELISA of Tsp-1 expression (normalized to total protein levels) in PC3M-LN4 fractions eluted from a Heparin-sepharose-Cu^2+^ column; **(E)** Western blot of PRSS2 and actin protein levels in PC3 and PC3M-LN4 prostate cancer cells; **(F)** Western blot of PRSS2 and actin protein levels in MDA-MB-231 (231), MDA-MB-231-LM2 (LM2), MCF7 and SUM159 breast cancer cells; **(G)** Tsp-1 and actin levels in WI-38 cells that were untreated (-) or treated with recombinant human PRSS2 (+); **(H)** Upper panel: Western blot of PRSS2 in SUM159 cells that were untransfected (C), transfected with empty pLKO.1 vector (V), or pLKO.1 vector expressing 3 independent shRNA sequences directed against PRSS2 (sh1, sh2, sh3); Lower Panel: Western blot of Tsp-1 and actin in WI-38 cells that were untreated (-) or treated with conditioned media from SUM159 cells with empty vector (V) or PRSS2 shRNA; **(I)** Western blot of Myc, PRSS2 and actin in SUM159 cells transfected with vector control (V) or three shRNA sequences directed against c-Myc (sh1, sh2 and sh3); **(J)** Western blot of Myc, PRSS2 and actin in PC3M-LN4 cells transfected with vector control (V) or three shRNA sequences directed against c-Myc (sh1, sh2 and sh3);

To identify the tumor-secreted protein responsible for the paracrine repression of Tsp-1 we utilized a proteomic screening method previously used to identify prosaposin as a stimulator of Tsp-1 expression (Figure 1C) ^5^. Briefly, CM was fractionated over a heparin-sepharose/Cu^2+^ column and proteins were eluted with a linear gradient of NaCl plus 10 mM imidazole. Collected fractions were dialyzed into PBS and used to treat primary human lung fibroblasts and Tsp-1 expression was analyzed by ELISA and western blot. We found that the Tsp-1 repressing activity was present in fractions that eluted with 1.0 and 1.1 M NaCl (Figure 1D and Supplemental Figure S1). These two fractions, along with the inactive fractions that eluted at 0.9 and 1.2M NaCl, were concentrated and analyzed by tandem liquid chromatography-mass spectrometry (LC-MS) analysis (Supplemental Figure S2). Analysis of the LC-MS results yielded only one protein that was present in both of the Tsp-1 repressing fractions and absent in the inactive fractions, the serine protease PRSS2. PRSS2 is an anionic trypsinogen and is also referred to as tumor-associated trypsin (TAT). PRSS2 has been found at elevated levels in tissue and serum of gastric, pancreatic, prostate and ovarian cancer patients ^22–26^.

We validated expression of PRSS2 in PC3M-LN4 cells by western blot and found that PC3M-LN4 cells expressed ~10-fold higher levels of PRSS2 than the weakly metastatic PC3 cells, which stimulate Tsp-1 expression in lung fibroblasts ^5^ (Figure 1E). We then analyzed a set of human breast cancer cell lines for PRSS2 expression and found that its expression level correlated with the metastatic potential of the cell lines (Figure 1F). Specifically, MDA-LM2, a metastatic derivative of MDA-MB-231 expressed significantly higher levels of PRSS2 than its parental cell line. Additionally, SUM159, a metastatic triple negative breast cancer cell line expresses significantly higher levels of PRSS2 than the ER+ and non-metastatic cell line MCF7.

### PRSS2 is necessary and sufficient for paracrine repression of Tsp-1

Having identified PRSS2 as being present in Tsp-1 repressing fractions of PC3M-LN4 cells we sought to validate that it represses Tsp-1. We treated primary human lung fibroblasts with purified recombinant human PRSS2 (rhPRSS2) and found that rhPRSS2 was sufficient to repress Tsp-1 expression (Figure 1G).

To determine if PRSS2 was required for the repression of Tsp-1 we silenced its expression in the highly metastatic breast cancer cell line SUM159 via lentiviral transduction of shRNA specific for PRSS2 (Figure 1H). Consistent with the proposed activity of PRSS2, we found that the level of Tsp-1 induction in target cells correlated with the level of repression of PRSS2 with three independent shRNA constructs transduced into SUM159 cells (Figure 1H). These findings indicate that PRSS2 is necessary and sufficient for the repression of Tsp-1 in human lung fibroblasts.

We previously demonstrated that the expression of Tsp-1 and prosaposin is repressed in a c-Myc-dependent manner in highly metastatic cells PC3M-LN4 (derived from PC3 cells) and MDA-LM2 (derived from MDA-MB-231 cells) ^5^. Due to the observation that these metastatic cell lines repressed prosaposin concomitantly with the upregulation of PRSS2, we examined whether PRSS2 expression was also regulated by c-Myc. shRNA-mediated silencing of c-Myc resulted in a reduction of PRSS2 expression in SUM159 (Figure 1I) and PC3-LN4 (Figure 1J) cells consistent with the level of c-Myc knockdown. These findings indicate that PRSS2 expression is c-Myc dependent and regulated in the opposite manner as prosaposin.

### PRSS2 enzymatic activity is not required for repression of Tsp-1

The observation that PRSS2 expression is necessary and sufficient to repress Tsp-1 in a paracrine acting fashion, suggests two possible mechanisms. The first is that PRSS2 acts as a protease to cleave a substrate that then interacts with a cell surface receptor to repress Tsp-1. The second is that PRSS2, itself, is a ligand for a cell surface receptor and represses Tsp-1 by direct binding and activation of a signal transduction cascade culminating in the repression of Tsp-1.

To test the first hypothesis, we generated point mutations in PRSS2 that inactivate its enzymatic activity. In addition to making mutations in the active site, by mutating the serine at residue 200 to alanine, threonine and cysteine (S200A, S200T, and S200C) we also generated a glycine to arginine substitution at residue 191 (G191R). This mutation has been identified as an inactivating mutation that confers resistance to familial pancreatitis^27^. We then ectopically expressed these four mutant versions of PRSS2 in 293T cells and used the conditioned media to treat lung fibroblasts. We found that all of the mutant proteins were expressed at levels comparable to wild-type PRSS2 protein (Figure 2A). We then used a colorimetric assay to measure the enzymatic activity of the mutant PRSS2 proteins. The results of this assay confirmed that the G191R and S200A mutants were enzymatically inactive and S200T and S200C retained minimal enzymatic activity (Figure 2B). Strikingly, the CM containing the PRSS2 mutants was able to repress Tsp-1 to the same relative degree as the wild-type protein (Figure 2C). These findings indicate that the enzymatic activity of PRSS2 is not required for repression of Tsp-1 expression and suggest that PRSS2 may function as a ligand for a cell surface receptor.

**Figure 2.**
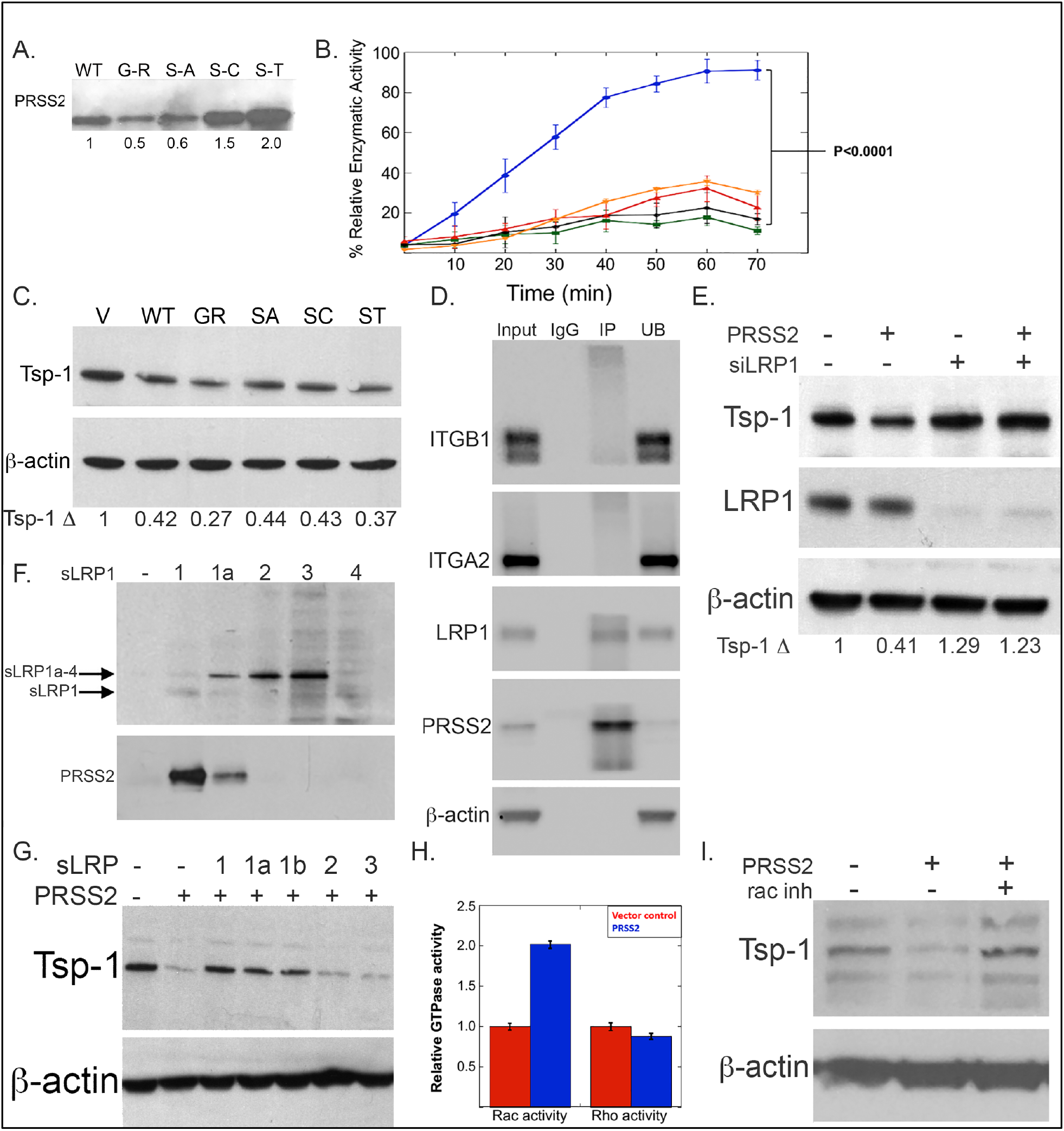
PRSS2 represses Tsp-1 via enzyme independent binding to LRP1. **(A)** Western blot expression of PRSS2 protein levels in conditioned media of 293T cells transfected with WT PRSS2 (blue), PRSS2-G191R (black), PRSS2-S200A (blue), PRSS2-S200C (red), and PRSS2-S200T (orange); **(B)** Plot of enzymatic activity WT and mutant PRSS2 proteins relative to WT PRSS2 (%); **(C)** Western blot of Tsp-1 and actin expression in WI38 fibroblasts treated with CM from 293T cells transfected with empty pCMV-Sport6 vector or vector expressing wild-type (WT) or a mutant version of PRSS2 (GR=G191R, SA=S200A, SC=S200C, ST=S200T); **(D)** Western blot of Integrin α2 (ITGA2), Integrin β1, ITGB1, LRP1, PRSS2, and actin following immunoprecipitation experiment of WI38 cells, using control IgG, or α-PRSS2 (IP) antibody and unbound protein (UB); **(E)** Western blot of Tsp-1, LRP1 and actin in WI38 cells that were untreated (-) or treated with PRSS2 (+) in the presence (+) or absence (-) of siRNA directed against LRP1; **(F)** Western blot of secreted truncation mutants of LRP1 (sLRP1) containing binding domains 1-4, PRSS2, and actin from an immunoprecipitation experiment with α-myc epitope antibody; **(G)** Western blot of Tsp-1 and actin in WI38 cells that were untreated (-) or treated with PRSS2 (+) in the absence (-) or presence of secreted truncated mutants of PRSS2 comprised of binding domains 1, 1a, 1b, 2 and 3; **(H)** Plot of GTP-bound Rac and Rho in WI38 cells treated with CM from 293T cells transfected with empty vector (red bars) or PRSS2 (blue bars); **(I)** Western blot of Tsp-1 and actin expression in WI38 cells that were untreated (-) or treated with PRSS2 (+) in the absence (-) and presence (+) of a small molecule inhibitor of Rac1.

### PRSS2 is a novel ligand for LRP-1

An examination of the existing literature revealed no reports of PRSS2 as a ligand for a cell surface receptor. To identify potential PRSS2 interacting proteins we performed a coimmunoprecipitation experiment, in which CM from 293T cells transfected with a plasmid expressing PRSS2 was mixed with cell lysates from WI-38 lung fibroblasts and PRSS2 was immunoprecipitated. The immunoprecipitates were run on an SDS-polyacrylamide gel, visualized by silver staining (Supplementary Figure S3) and subsequently analyzed by LC/MS. Results of the LC/MS analysis identified 325 proteins, with a minimum of three peptide fragments, that were present in the immunoprecipitate of α-PRSS2 but not the IgG control (Supplemental Figure S4). We performed gene ontogeny analysis (Panther) and independently searched the list of PRSS2-precipitated proteins for known cell surface receptors. This analysis yielded only 3 potential candidates: integrin α2 (ITGA2), integrin β1 (ITGB1), and low density lipoprotein-related receptor protein 1 (LRP1).

To confirm that PRSS2 specifically binds these proteins we repeated the immunoprecipitation experiment and performed western blot analysis for each receptor. We found that PRSS2 reproducibly co-immunoprecipitated with LRP1 but was not able to co-immunoprecipitate ITGA2 or ITGB1 (Figure 2D). To functionally validate these findings, we silenced expression of LRP1 in primary lung fibroblasts via siRNA and confirmed knockdown by western blot (Figure 2E). We then treated these cells with conditioned media from 293T cells that were transiently transfected with PRSS2 and observed that PRSS2 was unable to repress Tsp-1 in cells in which LRP1 expression was silenced. From these results we concluded that LRP1 is the cell surface receptor that mediates PRSS2 repression of Tsp-1 (Figure 2E).

As this is the first observation of PRSS2 binding to LRP1 we sought to identify the binding domain of LRP1 that mediates the interaction between the two proteins. For this experiment we incubated the CM from SUM159 cells with CM from 293T cells transfected with vectors expressing soluble (secreted) truncation mutants of LRP1 consisting of the four binding domains of the protein fused to an N-terminal Myc epitope tag ^28^. We found that only LRP1 binding domain 1 co-immunoprecipitated with PRSS2 (Figure 2F). While LRP1 has been reported to have over 50 ligands^29^, PRSS2 represents the first identified protein to bind independently to LRP1 domain 1 (as opposed to proteins, such as α2-macroglobulin, which bind to domain 1 but not independently of binding to domain 2 and 4)^28–30^.

We then tested the functional significance of the interactions between PRSS2 and LRP1 by treating lung fibroblasts with CM from 293T cells ectopically expressing PRSS2 in the presence and absence of soluble versions of the LRP1 truncation proteins. The rationale behind this experiment is that if the interactions between PRSS2 and domains 1 and 3 of LRP1 are required for the repression of Tsp-1, then the soluble truncated proteins should serve as decoys to sequester PRSS2 and prevent it from binding to LRP1. Consistent with the results from the co-IP experiment, incubation of PRSS2 with binding domains 1 and 1a of LRP1 blocked the repression of Tsp-1, while incubation with binding domains 2 and 3 had no effect (Figure 2G).

Finally, we sought to determine the components of the signal transduction cascade downstream of LRP1 leading to the repression of Tsp-1. It has been demonstrated that LRP1 ligands can activate Rho and Rac GTPases^31,32^. To determine whether Rho or Rac was activated by PRSS2 binding to LRP1 we measured the activation of both GTPases using plate-based G-LISA assays. We found that CM from 293T cells ectopically expressing PRSS2 stimulated Rac-GTPase activity ~2-fold in WI-38 cells but had no effect on Rho-GTPase activity (Figure 2H).

To functionally validate these findings we treated lung fibroblasts with CM from SUM159 cells in the presence and absence of a small molecule inhibitor of Rac1. We found that while CM from SUM159 cells repressed Tsp-1 by 50%, treatment with the Rac1 inhibitor in combination with SUM159 CM resulted in Tsp-1 levels that were 2.6-fold higher than CM alone and 1.3-fold higher than untreated cells (Figure 2I). These findings demonstrate that PRSS2 represses Tsp-1 via activation of Rac1 downstream of binding to LRP1.

Since LRP1 is also an endocytic receptor, we sought to determine whether PRSS2 was taken up by fibroblasts following treatment with CM from PRSS2-overexpressing 293T cells. We found that wild-type fibroblasts had increased levels of PRSS2 within one hour of treatment, but cells silenced for LRP1 did not (Supplemental Figure S5). These findings suggest that in addition to inducing a Rac-mediated signal transduction cascade leading to the repression of Tsp-1, PRSS2 is also endocytosed by LRP1.

### PRSS2 levels correlate with aggressive features of breast and prostate carcinoma

To determine whether PRSS2 has potential clinical relevance as a potential therapeutic target, we evaluated its expression by immunohistochemistry in series of human breast and prostate cancers (Figure 3A and B). In a population-based breast cancer series (Series 1, n=518), strong and consistent associations were found between PRSS2 expression and multiple features of aggressive tumors, such as high histologic grade, lack of expression of estrogen receptor (ER), tumor cell proliferation (by Ki67 expression), CK5/6 expression (basal marker), and increased angiogenesis (by pMVD and GMP) (Table 1). In a *BRCA-based* breast cancer series (Series 2, n=202), which was enriched for cases with germline *BRCA1* or *BRCA2* mutations, strong associations with high histologic grade, tumor cell proliferation (by mitotic count), and p53 expression were present (Supplementary Table 1). In contrast, no associations with *BRCA*-status were found. Notably, these associations (Series 1-2) were found for PRSS2 levels in both tumor epithelium and when recorded separately in the stromal compartment. PRSS2 levels in tumor cells and in the associated stroma was significantly associated (Spearman’s correlation: Series 1, rho=0.36, p<0.0005; Series 2, rho=0.45, p<0.0005).

**Figure 3.**
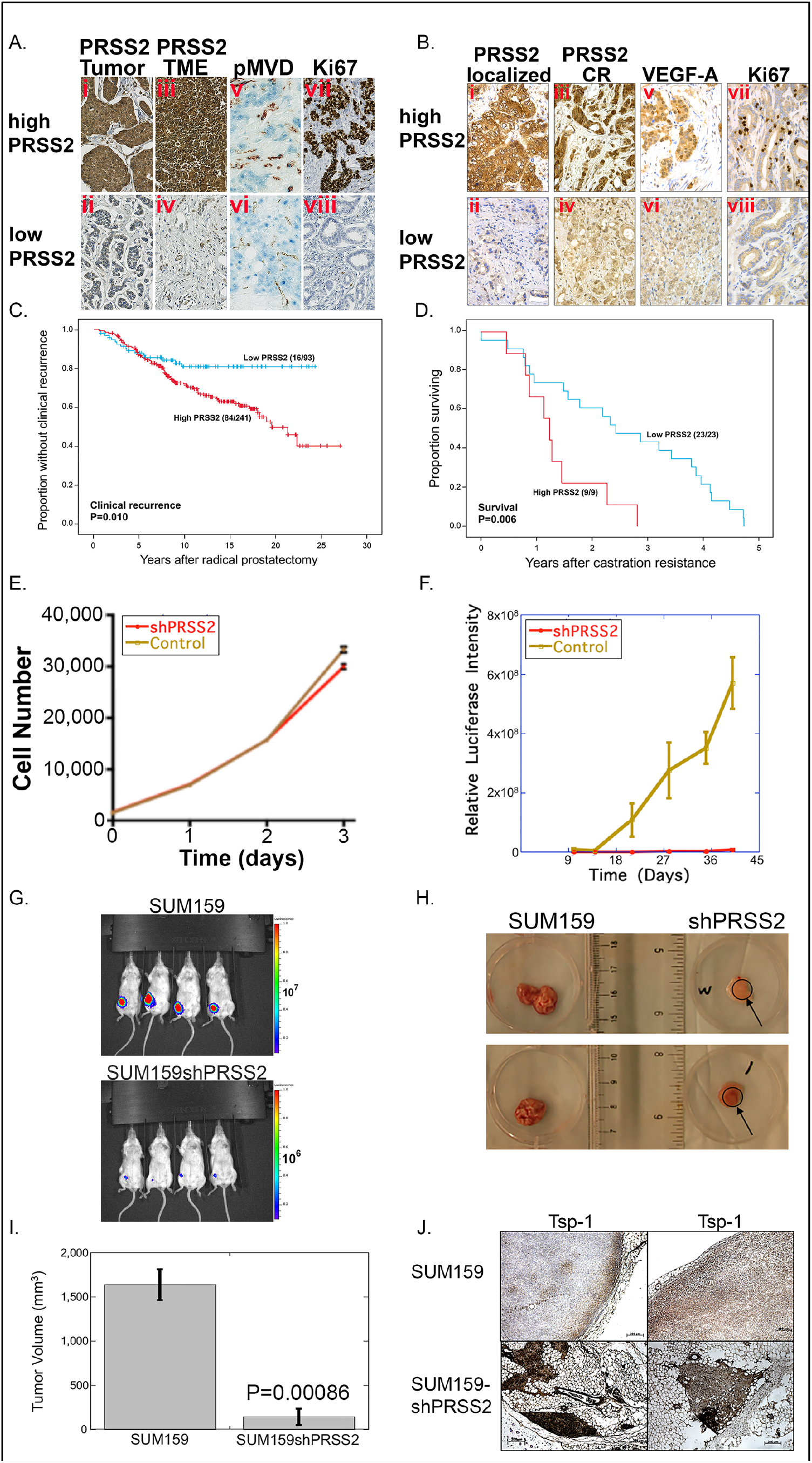
PRSS2 is required for efficient tumor formation of SUM159 cells. **(A)** Immunohistochemical analysis of breast cancer patient Series 1 for expression of: (**i**) PRSS2 in tumors with strong PRSS2 expression in tumor cells (high), (**ii**) PRSS2 in tumors with weak (low) PRSS2 expression in tumor cells, (iii) PRSS2 in tumors with strong PRSS2 expression in TME, (**iv**) PRSS2 in tumors with weak PRSS2 expression in the TME; **(v)** proliferating microvessel density (pMVD) in tumors with strong PRSS2 expression, (**vi**) pMVD in tumors with weak PRSS2 expression; (vii) tumor cell proliferation (Ki67) in tumors with strong PRSS2 expression; and (**viii**) Ki67 in tumors with weak PRSS2 expression. Original magnification x400. (**B**) Immunohistochemical analysis of prostate cancer patient series for expression of: (**i**) PRSS2 in tumors with strong (high) tumor cell expression of PRSS2 in localized prostatic carcinoma; (**ii**): PRSS2 in tumors with weak (low) tumor cell expression of PRSS2 in localized prostatic carcinoma; (**iii**) PRSS2 in tumors with strong expression of PRSS2 in castration resistant carcinoma (CR); (**iv**) PRSS2 in tumors with weak expression of PRSS2 in castration resistant carcinoma; (**v**) VEGF-A in localized carcinoma with strong PRSS2 expression; (**vi**) VEGF-A in localized carcinoma with weak PRSS2 expression; (**vii**) Ki67 in localized carcinoma with strong PRSS2 expression; (**viii**) Ki67 in localized carcinoma with weak PRSS2 expression. Original magnification x400. (**C**) Kaplan-Meier curve of clinical progression following radical prostatectomy of prostate cancer patients with strong and weak expression of PRSS2; (**D**) Kaplan-Meier curve of overall survival following the acquisition of castration resistance of prostate cancer patients with strong and weak expression of PRSS2; **(E)** Plot of *in vitro* proliferation of SUM159 and SUM159shPRSS2 cells over 3 days; **(F)** Plot of *in vivo* luciferase activity of orthotopic tumors formed by mammary gland injection of SUM159 and SUM159shPRSS2 cells; **(G)** Pictures of *in vivo* luciferase imaging of mice bearing tumors formed by SUM159 (upper panel) and SUM159shPRSS2 (lower panel) cells; **(H)** Photographs of mammary tumors formed by SUM159 and SUM159shPRSS2 cells; **(I)** Plot of volume of tumors formed by SUM159 and SUM159shPRSS2 cells; **(J)** Immunohistochemistry of Tsp-1 in tumors formed by SUM159 and SUM159shPRSS2 cells (scale bar=200μm).

**Table 1.**
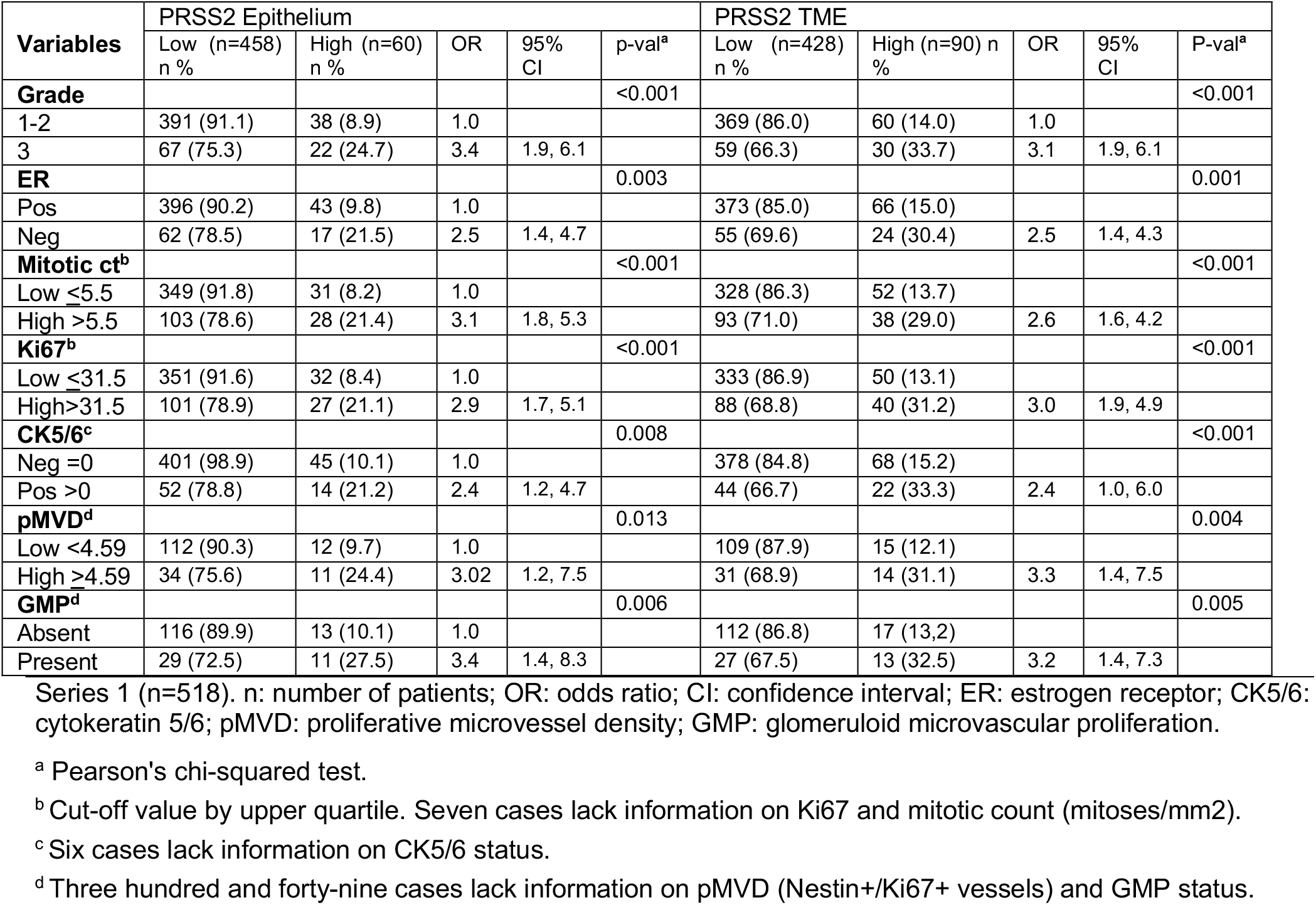
Breast Cancer Series 1.

In localized prostate cancer (Series 1, n=338), PRSS2 expression in tumor cells was associated with increased tumor cell proliferation (by Ki67 expression), and with increased VEGF-A (Figure 3B and Table 2). By univariate survival analyses (Series 1; n=338), strong PRSS2 was associated with shorter time to clinical recurrence (P=0.010), and it was borderline associated with biochemical recurrence (P=0.071) (Figure 3C and Supplementary Figure S6). By multivariate survival analyses (Series 1, n=338), where PRSS2 was included in addition to the standard prognostic variables Gleason score (≥4+3 versus ≤3+4), pathological stage (≥pT3 versus pT2) and preoperative s-PSA (dichotomized by upper quartile), strong PRSS2 independently predicted biochemical recurrence, loco-regional recurrence, and clinical recurrence (HR=1.4-2.5, P= 0.001-0.052), together with Gleason score and pathological stage, and for biochemical recurrence, s-PSA.

**Table 2.**
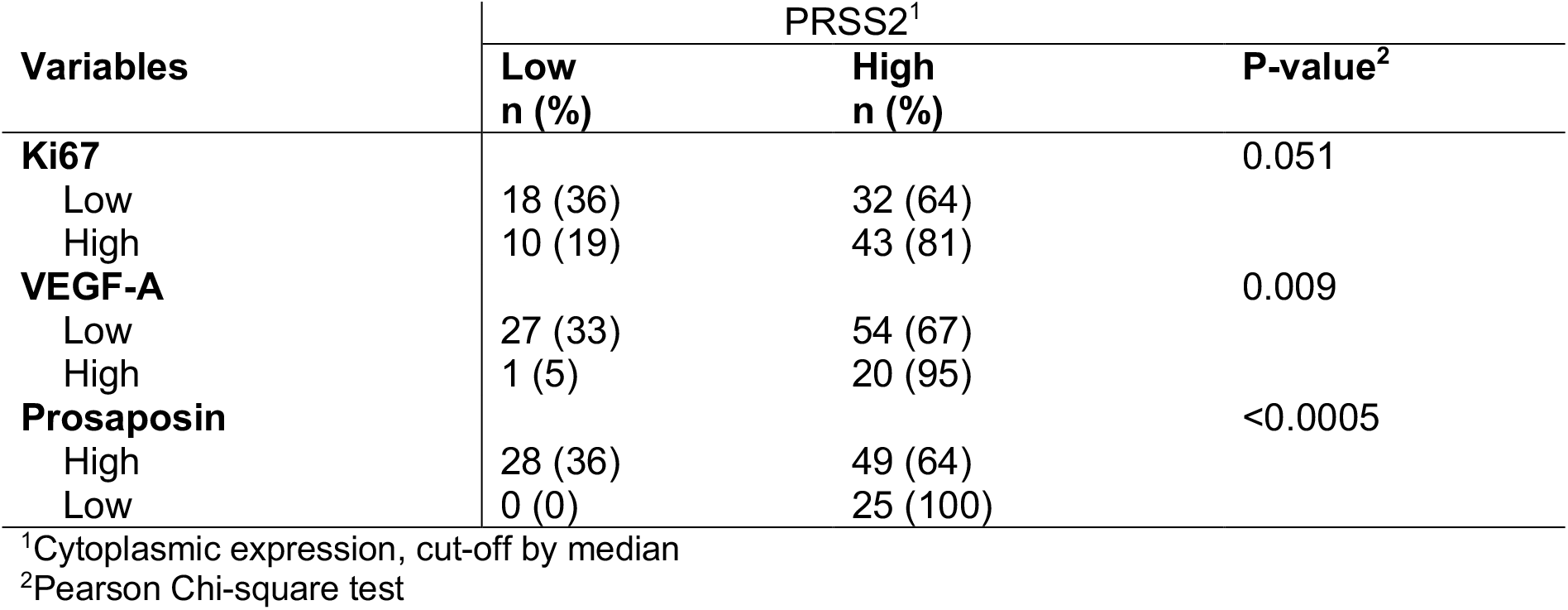
Associations between PRSS2 expression and selected biomarkers in localized prostatic carcinomas (radical prostatectomies)

| Variables | PRSS2<sup>1</sup> |  | P-value<sup>2</sup> |
| --- | --- | --- | --- |
|  | Low<br>n (%) | High<br>n (%) |  |
| **Ki67** |  |  | 0.051 |
| Low | 18 (36) | 32 (64) |  |
| High | 10 (19) | 43 (81) |  |
| **VEGF-A** |  |  | 0.009 |
| Low | 27 (33) | 54 (67) |  |
| High | 1 (5) | 20 (95) |  |
| **Prosaposin** |  |  | <0.0005 |
| High | 28 (36) | 49 (64) |  |
| Low | 0 (0) | 25 (100) |  |

<sup>1</sup>Cytoplasmic expression, cut-off by median
<sup>2</sup>Pearson Chi-square test

Strikingly, and consistent with the observation that c-Myc oppositely regulates PRSS2 and PSAP, PRSS2 was strongly and inversely associated with PSAP expression (**Table 2**). Specifically, of the 25 patients with low levels of PSAP, all had high levels of PRSS2. Finally, by univariate survival analyses of castration resistance prostate cancer (n=32), strong PRSS2 expression was associated with shorter time from diagnosis to death (P=0.006) (Figure 3D).

### PRSS2 is required for efficient tumor growth

The observations that more aggressive breast and prostate cancer cell lines express higher levels of PRSS2 than their less metastatic counterparts coupled with the corroboration of these findings in patient samples led us to hypothesize that disrupting PRSS2 expression could inhibit tumor growth. To test this hypothesis we silenced PRSS2 in SUM159 cells via shRNA and measured the proliferation rates of vector control and shPRSS2 cells. We found that vector control and SUM159 shPRSS2 cells had no statistically significant difference in *in vitro* cellular proliferation (Figure 3E). We then injected 1×10^6^ vector control and SUM159shPRSS2 cells, expressing firefly luciferase, orthotopically into the mammary gland of SCID mice (n=8 mice per cohort). After 40 days, the vector control tumors had reached a significant size, as determined by luciferase intensity and gross visual inspection (Figure 3F-I). Conversely, the SUM159-shPRSS2 cells formed tumors that were barely detectable by luciferase activity and undetectable by gross visual inspection (Figure 3F-I). In fact, 5/8 mice injected with SUM159shPRSS2 cells developed tumors smaller than 4 mm^3^ (less than 2 mm in diameter) and on average the tumors were 11.5-fold smaller than tumors formed by SUM159 vector control cells (1657 mm^3^ vs 144 mm^3^, P<0.001 by Mann-Whitney U test) (Figure 3F and I).

We then examined the tumors histologically by H&E and IHC for Tsp-1 expression. We found that Tsp-1 expression was significantly higher in tumors formed by the shPRSS2 cells than the vector control cells, where it was almost undetectable (Figure 3J). Of note, the size of the shPRSS2 tumors (1-2 mm in diameter) was consistent with previously published reports of dormant tumors in which Tsp-1 was highly expressed, either endogenously or ectopically ^33,34^. These findings suggest that tumor-secreted PRSS2 stimulates tumor growth via paracrine signaling to repress Tsp-1 in the tumor microenvironment and not via tumor cell autonomous effects.

### Loss of PRSS2 does not inhibit tumor growth in the absence of Tsp-1

Based on the observation that silencing PRSS2 in SUM159 cells significantly inhibited tumor growth, we asked whether this was due to the inability to repress Tsp-1 in the TME. We postulated that in the absence of Tsp-1, silencing PRSS2 should have little to no effect on tumor growth. To test this hypothesis, we made use of Tsp-1^-/-^ mice and silenced PRSS2 in the syngeneic C57B6/J derived pancreatic cancer cell line, Pan02 ^35^ (Supplemental Figure S7). We examined the effect of PRSS2 in pancreatic tumor growth and progression due to the reports that PRSS2 overexpression is a contributing factor to pancreatitis, which is a precursor lesion for pancreatic cancer ^36,37^. Specifically, we injected 5×10^5^ vector control Pan02 and Pan02shPRSS2 cells orthotopically into the head of the pancreas of wt and Tsp-1^-/-^ C57Bl6/J mice (n=8 mice per cohort) (Figure 4A). We monitored tumor growth via in vivo luciferase imaging and euthanized all mice when the wt mice injected with vector control Pan02 cells became moribund due to ascites development at 28 days post-injection. We then measured the volume and mass of all tumors and found that in wt mice, the tumors formed by Pan02shPRSS2 cells were <50% the size of vector control Pan02 tumors (528 mg vs 1082 mg; P=0.002, by Mann-Whitney U test) (Figure 4B and C). Conversely, in Tsp-1^-/-^ mice, the difference in the size of tumors formed by Pan02shPRSS2 cells compared to tumors formed by vector control Pan02 cells was not statistically significant (711 mg vs 779 mg; P=0.588 by Mann-Whitney U test (Figure 4C). We also stained all tumors for Tsp-1 expression via immunohistochemistry and found that in wt mice, vector control Pan02 tumors had little to no detectable Tsp-1 expression, while Pan02shPRSS2 tumors had significantly higher Tsp-1 levels (Figure 4D). As expected, all of the tumors in Tsp-1^-/-^ mice had no detectable Tsp-1 by IHC (Figure 4D).

**Figure 4.**
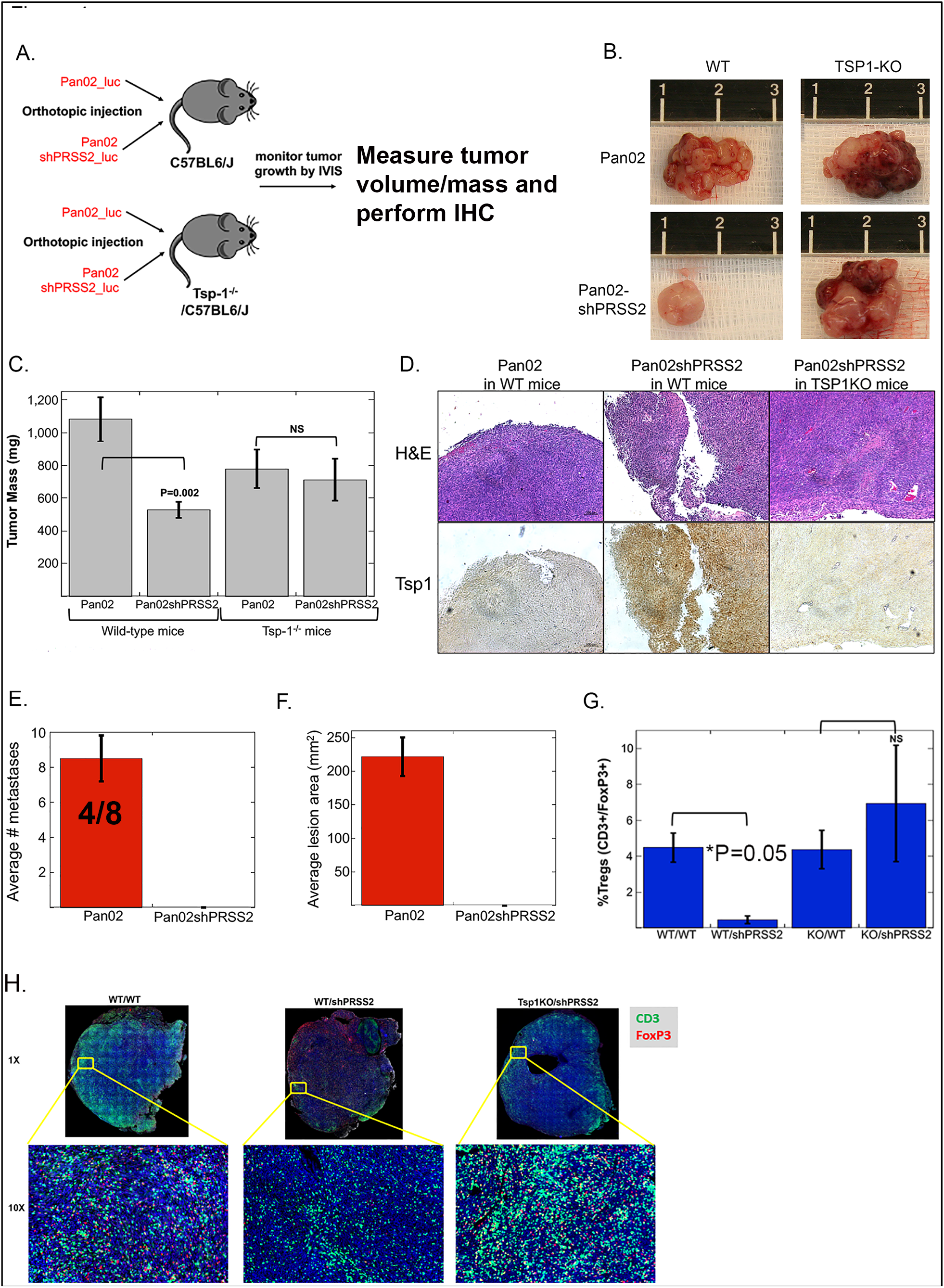
Silencing PRSS2 inhibits primary pancreatic tumor growth and metastasis. **(A)** Schematic of tumor implantation strategy; **(B)** Photographs of tumors formed by Pan02 and Pan02shPRSS2 cells in wild-type C57Bl6/J mice and *thbs1*^-/-^ C57Bl6/J mice; **(C)** Plot of average mass of tumors formed by Pan02 and Pan02shPRSS2 cells in wildtype C57Bl6/J mice and *thbs1*^-/-^ C57Bl6/J mice; **(D)** H&E and Tsp-1 immunohistochemical staining of tumors formed by Pan02 and Pan02shPRSS2 cells in wild-type C57Bl6/J mice and *thbs1*^-/-^ C57Bl6/J mice (Scale bar=200μm); **(E)** Plot of the average number of macrometastatic lesions identified by gross examination of the peritoneal cavities of mice bearing tumors formed by Pan02 and Pan02shPRSS2 cells in wild-type C57Bl6/J mice (4/7 refers to the number of mice that developed macrometastatic lesions); **(F)** Plot of the average area of macrometastatic lesions (in mm^2^) identified by gross examination of the peritoneal cavities of mice bearing tumors formed by Pan02 and Pan02shPRSS2 cells in wild-type C57Bl6/J mice; **(G)** Plot of percentage of CD3+/FoxP3+ T cells out of total CD3+ T cells in in tumors formed by Pan02 and Pan02shPRSS2 cells in WT C57Bl6/J mice and *thbs1*^-/-^ C57Bl6/J mice; and, **(H)** Immunofluorescence staining of CD3 (green) and FoxP3 (red) in tumors formed by Pan02 and Pan02shPRSS2 cells in WT C57Bl6/J mice and *Tsp1*^-/-^ C57Bl6/J mice.

Finally, upon gross examination of mice at the time of necropsy, we observed that 4/8 mice injected with Pan02 vector control cells developed large metastases on the wall of the peritoneal cavity and diaphragm, while 0/8 mice injected with Pan02shPRSS2 cells developed detectable metastases (Figure 4E and F). These findings demonstrate that loss of PRSS2 significantly inhibits primary tumor growth and metastasis. However, in the absence of Tsp-1, loss of PRSS2 does not inhibit tumor growth, indicating that the major role of PRSS2 in tumor growth is the paracrine-mediated repression of Tsp-1.

### Repression of Tsp-1 creates an immunosuppressive tumor microenvironment

Given the profound effects of silencing PRSS2 on tumor growth and progression we sought to determine whether modulating Tsp-1 expression in the TME had any effect on the tumor immune landscape. It has been demonstrated that regulatory T cells (T regs) express high levels of CD36, which makes them metabolically dependent on free fatty acid uptake ^38^. Tsp-1 binds to CD36 and blocks fatty acid uptake, in addition to inducing apoptosis ^39,40^. Thus, we postulated that tumors in which PRSS2 was silenced, and consequently express high levels of Tsp-1, would have fewer Tregs and a higher ratio of CD8+ T cells to Tregs. Accordingly, we analyzed Pan02 and Pan02shPRSS2 tumors that formed in WT and Tsp-1^-/-^ mice for T cell markers.

We performed immunohistochemical analysis of total T cells (CD3+), CD4+ T cells, CD8+ T cells, and FoxP3+ Tregs. In WT mice, we observed a 9-fold decrease of Tregs in Pan02shPRSS2 tumors compared to control Pan02 tumors (4.5% to 0.48% of all T cells; P=0.05, by Mann-Whitney U test) (Figure 4H and I). Strikingly, in Tsp-1^-/-^ mice the percentage of Tregs did not differ significantly between Pan02 and Pan02shPRSS2 tumors (4.4% vs 6.95%, P=0.63, by Mann-Whitney U test). Moreover, the percentage of CD3+/CD8+ T cells in shPRSS2 tumors was >2-fold higher than in vector control tumors in wild type mice (3.36% vs 1.59%; P=0.057 by Mann-Whitney U test) but was not significantly different in Tsp-1^-/-^ mice (3.97% vs 5.1%, P=0.7 by Mann-Whitney U test) (Supplemental Figure S8). Thus, the ratio of CD8+:FoxP3+ T cells was 20-fold higher in tumors formed by Pan02shPRSS2 cells (3.59:0.48=7 vs 1.59:4.5=0.35). These findings indicate that the repression of Tsp-1 results in the generation of an immunosuppressive tumor microenvironment with high levels of regulatory T cells and a low ratio of CD8+:Treg cells. Our results indicate that blocking the repression of Tsp-1 dramatically increases CD8+ T cells and concomitantly reduces Treg infiltration which could result in a more active tumor immune microenvironment.

### Myeloid-specific genetic deletion of LRP1 prevents repression of Tsp-1 by PRSS2

We have previously demonstrated that genetic deletion of Tsp-1, in the entire mouse or specifically in bone-marrow derived cells, abrogates the ability of prosaposin (PSAP) to inhibit tumor growth ^4,5^. Accordingly, we specifically deleted LRP1 in myeloid derived cells by crossing LysM-Cre mice with LRP1^flox/flox^ mice (Supplemental Figure S9A). FACS analysis revealed that LRP1 is predominantly expressed by CD11b+ myeloid cells, as opposed to T (CD3+) and B (B220+) cells (Figure 5A and B). Importantly, myeloid specific knock out of LRP1 did not affect production of myeloid cells in these mice (Figure 5C and Supplemental Figure S10).

**Figure 5.**
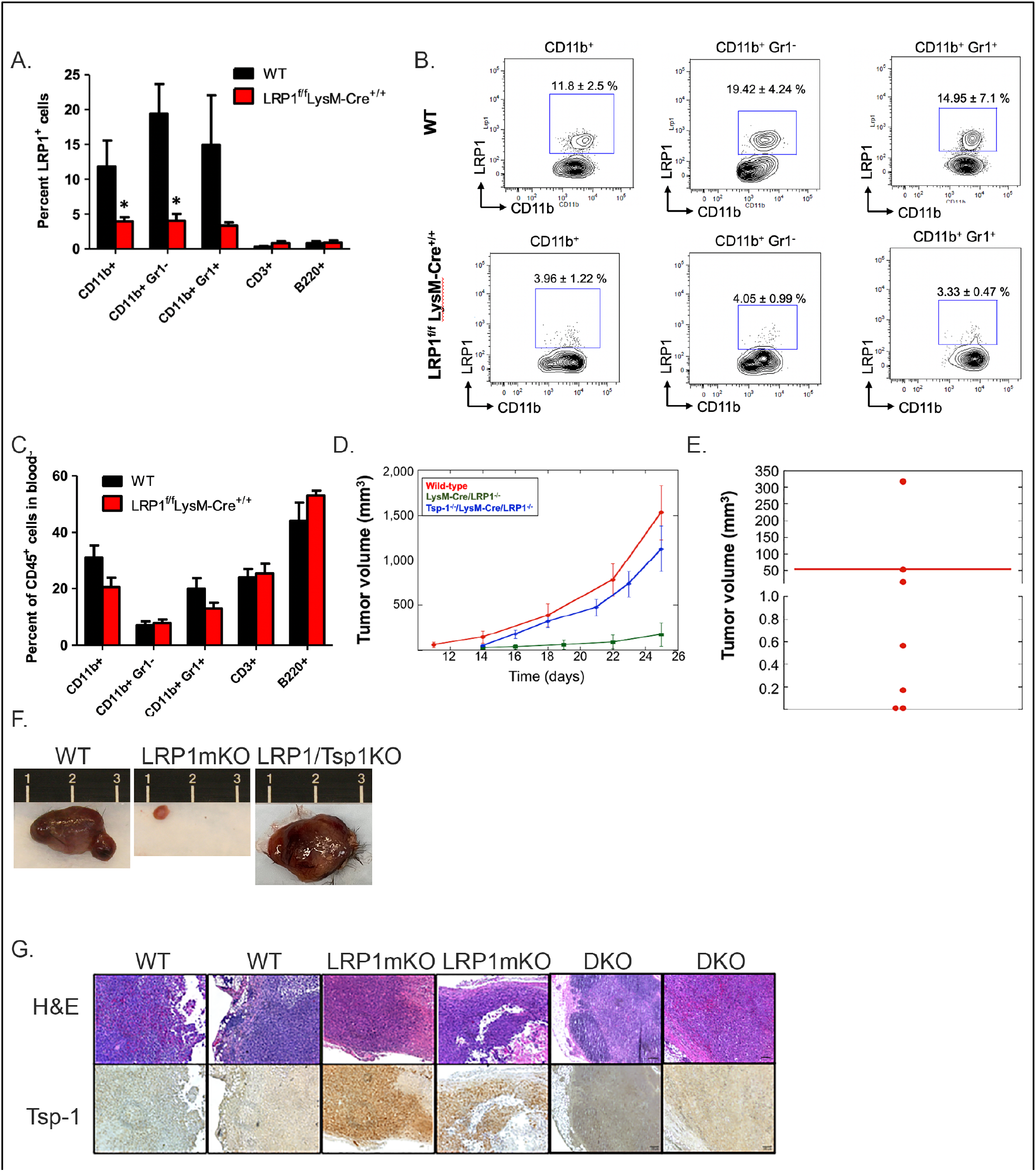
Myeloid specific deletion of LRP1 inhibits tumor growth by preventing the repression of Tsp-1. **(A)** Plot of average LRP1 expression in multiple myeloid and lymphoid cells in wild-type and LysM-Cre/LRP1^fl/fl^ mice as determined by FACS analysis; **(B)** Representative FACS plots of LRP1 expression in myeloid and lymphoid cells in wildtype and LysM-Cre/LRP1^fl/fl^ mice; **(C)** Plot of relative abundance of myeloid and lymphoid cells in wild-type and LysM-Cre/LRP1^fl/fl^ mice as determined by FACS analysis; **(D)** Plot of average tumor volume (as measured by calipers) of orthotopic mammary tumors formed by E0771 murine breast cancer cells in wild-type (red line), LysM-Cre/LRP1^fl/fl^ (green line), and LysM-Cre/LRP1^fl/fl^/THBS1^-/-^ (blue) mice; **(E)** Dot plot of volume of tumors formed by E0771 cells in LysM-Cre/LRP1^fl/fl^ mice; **(F)** Photographs of tumors formed by E0771 murine breast cancer cells in wild-type, LysM-Cre/LRP1^fl/fl^, and LysM-Cre/LRP1^fl/fl^/THBS1^-/-^ mice; **(G)** H&E and Tsp-1 immunohistochemical staining of tumors formed by E0771 murine breast cancer cells in wild-type, LysM-Cre/LRP1^fl/fl^, and Tsp-1^-/-^/LysM-Cre-LRP1^fl/fl^ mice (scale bars =100 μm).

We then sought to determine whether the disruption of PRSS2-LRP1 signaling in myeloid cells could inhibit primary tumor growth. We injected wild-type C57BL6/J mice and LysM-Cre-LRP1^-/-^ mice with the murine triple negative breast cancer cell line E0771. Strikingly, we found that tumors formed by E0771 tumors in the myeloid specific LRP1 knockout mice were 28.3-fold smaller than tumors formed in wild-type mice (54.9 mm^3^ vs 1,531 mm^3^; P=0.002 by Mann-Whitney U test) (Figure 5D and F). Moreover, 4 out of 8 tumors that formed in LysM-Cre-LRP1^-/-^ mice were less than 2 mm in diameter (Figure 5E). Histologic analysis of the tumors that formed in the LysM-Cre-LRP1^-/-^ mice revealed abundant Tsp-1 in the TME, while tumors in wt C57BL6/J mice had virtually undetectable levels of Tsp-1 (Figure 5G). To confirm that the loss of LRP1 inhibited tumor growth due to the inability of tumors to repress Tsp-1, we crossed the LysM-Cre-LRP1^-/-^ with Tsp1^-/-^ mice to generate double knockout (DKO) mice that lacked Tsp-1 globally and LRP1 in myeloid cells (Figure S9B). We injected 1×10^6^ E0771 cells into the mammary gland of the DKO mice and monitored tumor growth by direct measurement. We found that the average size of E0771 tumors that formed in the DKO mice ranged between 75-93% of the size of the tumors that formed in wild-type mice. Moreover, tumors that formed in the DKO mice had undetectable levels of Tsp-1 by IHC, as expected (Figure 5D-G). These findings indicate that LRP1 on myeloid cells is required for the PRSS2-mediated repression of Tsp-1 in the TME and that the inability to repress Tsp-1 in the TME significantly impairs tumor growth.

## Discussion

The ability of tumor cells to modify their microenvironment to create a site that is permissive for growth is a key factor in distinguishing aggressive, metastatic tumors from localized lesions. It has been previously reported that the tumor cell autonomous repression of Tsp-1 is required for tumors to escape dormancy ^33,34^. Here we demonstrate that repression of Tsp-1 in the tumor microenvironment is also required for tumors to escape dormancy.

In this report we describe the identification of a novel stimulator of tumor growth and metastasis, the serine protease PRSS2. Of note, we demonstrate that the tumor growth promoting activity of this protein lies not in its enzymatic activity but in its ability to act as a ligand for the cell surface receptor LRP1. In doing so we also discovered a novel biologic activity for the LRP1/Rac pathway, the repression of the anti-angiogenic, anti-tumorigenic, and anti-inflammatory protein Tsp-1.

Silencing PRSS2 in tumor cells significantly attenuates tumor growth and metastasis. Additionally, silencing PRSS2 also resulted in a TME that is less immunosuppressive as evidenced by increased infiltration of CD8+ T cells and decreased infiltration of Tregs resulting in an increased ratio of CD8:Treg. Critically, these alterations in immune cell composition of the TME were Tsp-1-dependent as they were not observed in Tsp-1^-/-^ mice. Finally, specific genetic deletion of the receptor for PRSS2, LRP1, in myeloid cells recapitulates the tumor inhibitory effects of silencing PRSS2.

Critically, the experimental findings that PRSS2 is required for efficient tumor growth were validated by the observation that PRSS2 expression correlates with aggressive clinical features such as angiogenesis, tumor cell proliferation, disease progression and survival in breast and prostate cancer patients. We previously demonstrated that tumors that express high levels of prosaposin (PSAP) remain localized and metastasize with very low frequency due to the induction of Tsp-1 expression in the TME ^4,5,16^. The progression, or lack thereof, of tumors could thus potentially be explained as a competition between PRSS2 and PSAP. If tumors make more PSAP than PRSS2 they will grow more slowly and if tumors make more PRSS2 than PSAP they will progress more rapidly. This hypothesis was born out by the observations that PRSS2 expression negatively correlates with PSAP expression in prostate cancer.

Based on the findings presented in this report, we predict that therapeutic strategies, which augment PSAP activity and inhibit PRSS2 binding to LRP1 could hold tremendous therapeutic potential for cancer patients. Significantly, this strategy should have relatively few adverse effects, as the therapeutic agents would not have direct cytotoxic activity. Moreover, since these therapeutic agents would target biological mechanisms in genetically normal cells that comprise the tumor microenvironment they would be less likely to induce resistance and potentially be indication agnostic.

## Author contributions

Conception and design: S. Wang, L. Sui, and R.S. Watnick; development of methodology: S. Wang, L. Sui, D.R. Bielenberg, D. Ganguly, Tina El Rayes, O.J. Halvorsen, A. Zeimys, R.S. Watnick; acquisition of data (provided animals, acquired and managed patients, provided facilities, etc.): S. Wang, L. Sui, D. Ganguly, Tina El Rayes, Cecilie Askeland, Astrid Børretzen, Danielle Sim, Ole Johan Halvorsen, Gøril Knutsvik, Sura Aziz, R.S. Watnick; analysis and interpretation of data (e.g., statistical analysis, biostatistics, computational analysis): S. Wang, D. Ganguly, O.J. Halvorsen, L.A. Akslen, A. Ziemys, L. Sui, R.A. Brekken, R.S. Watnick; administrative, technical, or material support (i.e., reporting or organizing data, constructing databases): R.S. Watnick, W.D. Foulkes, L.A. Akslen, R.A. Brekken; study supervision: R.A. Brekken, R.S. Watnick.

## Acknowledgements

We would like to thank Dr. Dudley Strickland (University of Maryland) for the soluble LRP1 truncation mutants and Kristin Johnson for artwork. We also would like to thank Richard O. Hynes and Bruce Zetter for helpful discussions. RW was supported by NIH R01 CA13547, the Elsa U. Pardee Foundation and the Strike 3 Foundation.

## Experimental Methods

### Cell culture and siRNA transfection

WI-38 cells were cultured in Minimal Essential Media (MEM), containing 10% fetal bovine serum (FBS) and 1% penicillin/streptomycin at 37°C. Cells were seeded the day before the experimentation and were subjected to PRSS2 transient transfection at a density of 80% confluence using Lipofectamine 3000 according to manufacturer’s protocol (Invitrogen). For siRNA transfection, WI-38 cells were transiently transfected with 25 nM siLRP1 (Sigma) using Lipofectamine 3000 according to manufacturer’s protocol (ThermoFisher). Gene silencing was confirmed by immunoblotting 72 hours post transfection.

### Treatment of cells with chemical inhibitors and recombinant protein

WI-38 cells were seeded 24 hours before the treatment and synchronized for 2 hours in serum-free medium. Synchronized cells were treatment with 25μm Rac1 inhibitor (Millipore), or 25μM Nutlin3 (Sigma), or PRSS2 conditioned medium for 16 hours and subjected to immunoblot analysis.

### Western blot analysis

Western blot was prepared as described. The following primary antibodies were used: Tsp1 (rabbit pAb, Abcam), p53 (DO-1 mouse mAb, Santa Cruz), LRP1 (rabbit pAb, Cell Signaling Technologies), β-actin (mouse mAb AC-15, Abcam).

### Generation of PRSS2 mutants

Single mutation of PRSS2 in pCMV-Sport6 vector (Harvard Plasmid Repository) (G191R, S200A, S200C and S200T) were introduced by site directed mutagenesis using QuikChange Lightning Site-Directed Mutagenesis Kit according to manufacturer’s protocol (Agilent Technologies). To express PRSS2 WT and Mutants protein, transient transfections were carried out in HEK293 cells cultured in 6-well culture plate using Lipofectamine 3000 and 2μg of PRSS2 wild-type or mutations were used. The transfection medium was replaced with serum-free DMEM medium 48 hours after transfection and conditioned medium containing secreted PRSS2 WT and mutation protein was collected after additional 24 hours.

### PRSS2 enzymatic activity assay

Recombinant PRSS2 and mutation protein was overexpressed in HEK293 cells as described above. Cells were cultured in DMEM phenol red free medium to reduce background reading for fluorescence assay. Conditioned medium (100μl) was supplemented with 0.1M Tris-HCl (pH 8.0), 1mM CaCl_2_ (final concentrations). PRSS2 activity were determined with 21.5 μM Mca-RPKPVE-Nval-WRK(Dnp)-NH_2_ Fluorogenic MMP Substrate (R&D Systems) (final concentration) and incubated at 25°C. Activity were measured every 10 minutes for total 70 minutes and expressed as percentage of potential total enzymatic activity. Triplicate experiments were performed for each construct.

### Co-Immunoprecipitation

WI-38 cells form 15cm dishes were collected and lysed in lysis buffer (25 mM Tris (pH 7.2), 150 mM NaCl, 5 mM MgCl2, 0.5% NP-40, 1 mM DTT, 5% glycerol) containing fresh protease inhibitor and phosphatase inhibitor cocktail (ThermoFisher), followed by centrifugation to remove cellular debris. Lysates were pre-cleared by incubating with 20 μl protein A/G agarose suspension for 1 hour at 4°C. The pre-cleared lysates were incubated with 10 μg PRSS2 antibody (Abcam) overnight at 4°C, followed incubation with 60 μl protein A/G agarose suspension for 3 hours at 4°C. Agarose beads were then collected by centrifugation, washed 4 times with 50mM Tris, 150mM NaCl, and 0.1% Tween-20, and resuspended in 100 μl RIPA buffer containing 6X SDS sample buffer. Samples were boiled for 5 minutes and the supernatant were collected after centrifugation for mass spectrometry analysis.

### Rac1 and RhoA GTPase activation assay

Rac1 and RhoA GTPase actitation was measured by G-LISA Rac1 activation assay Kit and G-LISA RhoA activation Kit (Cytoskeleton), according to manufacturer’s protocol. Briefly, WI-38 cells were serum starved for 2 hours, and the 293T conditioned medium with pCMV (control vector) and pCMV_PRSS2 vector overexpression was added to the serum starved cells for 45 min. The cells are washed with cold PBS and lysed in ice-cold lysis buffer containing protease inhibitors. Protein concentrations were measured, and the same amount of protein was used under each condition. The cell lysates were incubated in Rac1 and RhoA assay wells along with blank control and positive control for 30 min at 4°C with agitation at 400rpm. The wells were washed with Wash Buffer followed by incubation with Antigen Presenting Buffer at room temperature. The primary and secondary antibodies were incubated at room temperature for 45 min. The HRP detection reagent were incubated in each well for exact 3 min and the luminescence signal were detected using a microplate luminescence reader.

### Animal breast cancer model

All animal work was conducted in accordance with a protocol approved by the Institutional Animal Care and Use Committee (IACUC) of Boston Children’s Hospital. Female SCID mice (n=8/ group) (6-8 weeks old) were purchased from Mass General Hospital. For orthotopic breast cancer cell injection, SUM159 vector_luc and SUM159 shPRSS2_luc cells were injected into the mammary fat pad of the female SCID mice (1×10^6^ cells/ 20μl). The tumor burden was monitored weekly by both bioluminescence imaging using Xenogen IVIS system and tumor size measurement by caliper.

Myeloid-specific LRP1 knock out mice were generated by crossing LysM-Cre mice with LRP1^flox/flox^ mice. Murine triple negative breast cancer cell line E0771 (1×10^6^ cells/ 20μl) were orthotopically injected into the mammary fat pad of the female LysM-Cre-LRP1^-/-^ mice (n=8/group) (6 weeks old) and wild-type C57BL6/J mice (n=8/group) (6 weeks old). The tumor size was measured twice a week by caliper.

LRP1 and Tsp-1 double knock out (DKO) mice were generated by crossing LysM-Cre-LRP1^-/-^ with Tsp1^-/-^ mice. The double knockout was confirmed by genotyping using the following primers:

Lrp1^flox^_F: CATACCCTCTTCAAACCCCTTCCTG,

Lrp1^flox^_R: GCAAGCTCTCCTGCTCAGACCTGGA,

Tsp1_F: GAGTTTGCTTGTGGTGAACGCTCAG,

Tsp1wt_R: AGGGCTATGTGGAATTAATATCGG,

Tsp1ko_R: TGCTGTCCATCTGCACGAGACTAG.

E0771 cells (1×10^6^ cells/ 20μl) were orthotopically injected into the mammary fat pad of the female DKO mice (n=10/group) (6 weeks old). The tumor size was measured twice a week by caliper.

### Animal pancreatic tumor model

Thrombospondin-1 deficient mice (Tsp-1^-/-^, n=8/ group) were purchased from the Jackson Laboratory (#006141) and maintained by mating Tsp-1^-/-^ males with Tsp-1^-/-^ females. C57BL6/J mice (n=8/ group) (8 weeks old) were purchased from the Jackson Laboratory (#000664) and used as Tsp-1^+/+^ controls. For orthotopic pancreatic cancer cell injection, pancreatic cell lines wtPan02_luciferase and Pan02shPRSS2_luciferase (5 x 10^5^ cells/ 10 μl), washed and harvested in HBSS and mixed 1:1 with Matrigel, were injected into the tail of the pancreas (20 μl total volume). The tumor burden was determined weekly by bioluminescence imaging using Xenogen IVIS system initiated 7 days post injection.

### Thrombospondin-1 immunohistochemistry

Tumor samples were fixed in 10% paraformaldehyde and subsequently paraffin-embedded for sectioning. The paraffin-sectioned slides were deparaffinized with xylene and rehydrated in decreasing concentration of ethanol to water. For Thrombosponsdin-1 staining, the antigen retrieval was performed with proteinase K (Roche Diagnostics) at a final concentration of 20 μg/ml in 0.2 M Tris pH 7.2 at 37°C for 25 min. The slides were then blocked with 2.5% goat serum (Vector Laboratories) for 30 min at room temperature. Slides were incubated overnight at 4°C with primary antibody rabbit anti-Thrombospondin-1 (ab226383, Abcam) The slides were then washed in PBS with 0.05% Tween (3 times for 5 min), followed by incubation with HRP conjugated secondary antibody (Vector Laboratories) for 30 min at room temperature. Slides were then incubated with DAB substrate (Vector Laboratories) followed by counterstain with Hematoxylin (Vector lab, H3401).

### T-cell Immunohistochemistry

Immunohistochemistry (IHC) was performed as previously described ^41^. Briefly, slides were warmed in a 60 °C oven for 10 min followed by deparaffinization and rehydration. Before antigen retrieval, slides were fixed in 10% neutral buffered formalin for 30 minutes. Antigen retrieval was performed in antigen retrieval buffer (10 mM Tris-HCl, 1 mM EDTA with 10% glycerol [pH 9]) at 110 °C for 17 min (~4-5ψ). Slides were then allowed to be cooled down to room temperature and were washed once with PBS. Tissue sections were blocked with 2.5% goat serum (Vector Laboratories, S-1012) for 30 min followed by incubation with primary antibody overnight: CD4 (1:1000; Abcam, ab183685), Foxp3 (1:200; R&D Systems, MAB8214). After washing, the slides were incubated with HRP conjugated secondary Antibody (ImmPRESS; Vector Laboratories, MP-7401) for 30 min on a shaker. For developing the fluorescence signal, TSA detection system (PerkinElmer) was used. We used OPAL 520, OPAL 570 and OPAL 690 fluorophores for staining the different markers. Multiplex staining was performed by stripping the previous antibody in 10 mM citrate buffer (pH 6.2) plus 10% glycerol at 110 °C for 2 min before probing with the next primary antibodies in the next two consecutive rounds: CD3 (1:2000; Thermo Fisher Scientific, PA1-29547), and CD8 (1:4000; Cell Signaling, 98941). Slides were counterstained with DAPI and then cover slipped using ProLong Gold mount (no. P36931; Life Technologies). Slides were scanned at 20X using the Zeiss AxioScan.Z1 (Whole Brain Microscopy Facility, UT Southwestern). The following channels were used to acquire images: DAPI, AF488 (for OPAL 520), AF555 (for OPAL 570), and AF660 (for OPAL 690).

### Breast cancer patient series

Two independent breast cancer series were immunohistochemically stained for PRSS2 protein. Series 1 is a population-based cohort of 544 primary breast carcinomas from the period 1996-2003, and Series 2 is a case-control series of 202 primary breast carcinomas (53 *BRCA1*, 45 *BRCA2* and 104 *BRCA* non-mutated) from the period 1986-2005, as previously described^42,43^. Twenty-six cases from Series 1 and 30 cases from Series 2 were excluded due to technical inadequate material, leaving 518 and 172 cases for evaluation of PRSS2 staining. Outcome data was only available for Series 1 and included survival time, survival status and cause of death. Last date of follow-up was June 30, 2017 (median follow-up time of survivors, 216 months; range 166-256). During follow-up, 87 patients (17%) died of breast cancer, and 199 (23%) died of other causes.

The study was approved by the Western Regional Committee for Medical and Health Research Ethics, REC West (REK 2014/1984) (Series 1) and the Institutional Review Board at McGill University Hospital, A03-M33-02A (Series 2). All studies were performed in accordance with guidelines and regulations by the University of Bergen and REK, and in accordance with the Declaration of Helsinki Principles.

### Clinico-pathologic variables

The following variables were available: age at diagnosis, tumor diameter, histologic type, histologic grade, lymph node status, hormonal (ER, PR) and HER2 receptor status, proliferation markers (Ki67, mitotic count), CK5/6 (basal marker), p53 protein expression, proliferating microvessel density (pMVD) (nestin and Ki67 co-expression, series 1; Factor VIII and Ki67 co-expression, series 2) ^44^. The glomeruloid microvascular phenotype (GMP), a marker of increased tumor associated angiogenesis, was available for a subset of Series 1 ^45^.

### PRSS2 Immunostaining

Manual staining for PRSS2 was primarily performed on tissue microarray (TMA) sections, and regular sections were used in cases with poor quality or insufficient tumor material for evaluation in the TMA cores (72 cases in Series 1). The sections (5 μm) with formalin-fixed and paraffin-embedded tissue were deparaffinized with xylene, rehydrated in decreasing concentrations of alcohol and rinsed in distilled water. The slides were boiled in buffered solution at pH6 (DAKO S1699) using a microwave oven for 20 min at 350 W. After 15 min of cooling and addition of distilled water to reduce the fluid to room temperature, the slides were moved to a humidifying chamber (Magnetic Immuno Staining Tray, Cell Path, UK). To reduce the endogenous peroxidase, a peroxidase-blocking agent (DAKO S2023) was added for 8 minutes. Between the different steps, rinsing with buffered saline solution (DAKO S3006) was performed. The sections were incubated for 60 minutes at room temperature with the rabbit antibody PRSS2 (Center) (Sigma-Aldrich, SAB 1307060) diluted 1:50 in antibody diluent with background reducing components (DAKO S 3022). A secondary antibody (HRP EnVision rabbit (DAKO K4003) was added for 30 minutes at room temperature. For visualization, DAB (DAKO K3468) was used as chromogen and the slides were counterstained with hematoxylin (DAKO S3301). Multiorgan TMA sections were included as positive and negative controls. The negative controls were obtained by adding antibody diluent without the primary antibody.

### Evaluation of PRSS2 staining in tumor cells and stroma

PRSS2 staining in tumor cells was recorded using a semi-quantitative and subjective grading system, considering the intensity of staining (none=0, weak=1, moderate=2 and strong=3) and the proportion of tumor cells showing a positive reaction (<10%=1, 10-50%=2, >50%=3). A staining index (values 0-9) was calculated as a product of staining intensity (0-3) and proportion of immunopositive cells (1-3) ^46^. The evaluation of PRSS2 staining in the tumor stroma (tumor microenvironment staining, TME) was a combined subjective recording of the intensity of staining in cells present in the stroma compartment, mainly immune cells and fibroblasts (none=0, weak=1, moderate=2 and strong=3). If the staining intensity in stromal cells was heterogenous, the scoring was based on the predominant pattern.

As there is yet no validated cut-off value for PRSS2 expression, the distribution and frequency histograms for SI and intensity were evaluated. PRSS2 expression in tumor cells was considered low for SI 0-6 (88%) and high for SI 9 (12%) in Series 1 (upper quartile), and low for SI 0-4 (66%) and high for SI 6-9 (34%) in Series 2 (median). Stromal PRSS2 expression was considered low for staining intensity 0-2 (83% and 76%), and high for staining intensity 3 (17% and 24%) in Series 1-2.

### Statistical analysis

Data were analyzed using the SPSS Statistics for Windows, Version 25.0. (IBM Corp., Armonk, NY, USA). Statistical significance was assessed at the two-sided 5% level, whereas borderline statistical significance was defined as P-values between 5% and 10%. Associations between categorical variables were evaluated using the Pearson’s χ^2^ test or Fisher’s exact test, as appropriate, and odds ratios (OR) were computed. Univariate survival analyses were carried out using the Kaplan-Meier method with significance determined by the log-rank test. The endpoint in survival analyses was breast cancer specific survival (BCSS) (Series 1). Entry date was the date of diagnosis. Patients who died from other causes were censored at the date of death.

### Prostate cancer patient series

Different series of prostatic tissues were used. Series 1 includes carcinoma tissues from 338 patients treated by radical prostatectomy for clinically localized prostate cancer (Haukeland University Hospital, Bergen, Norway, during 1986-2007), with follow-up until September 2016 (median follow-up for survivors 157.1 months; biochemical recurrence or clinical recurrence (local or distant) were recorded as endpoints). Series 2 includes tissues from 33 patients with castration-resistant prostate carcinoma receiving palliative treatment with transurethral resection of the prostate during 1990-2005 (follow-up with respect to deaths). Univariate survival analyses were carried out using the Kaplan-Meier method with significance determined by the log-rank test.

The study was approved by the Western Regional Committee for Medical and Health Research Ethics, REC West (REK 2015/2178) (Series 1,2). All studies were performed in accordance with guidelines and regulations by the University of Bergen and REK, and in accordance with the Declaration of Helsinki Principles.

### Clinico-pathologic variables

Clinico-pathologic information were retrieved from the clinical patient files and pathology reports for the patients in Series 1. The information included age at diagnosis, preoperative and postoperative s-PSA, clinical TNM stage, Gleason grading, largest tumor dimension, involvement of surgical margins, extra-prostatic extension, seminal vesicle invasion, and pelvic lymph node status at prostatectomy.

Follow-up information involved time from surgery until biochemical recurrence, clinical recurrence, loco-regional recurrence, skeletal metastasis and death, including prostate cancer specific death for the patients in Series 1, and time from castration resistance to death for the patients in Series 2.

Tissue microarrays consisted of three tissue cores from each case (diameter 0.6-1.0 mm), selected from areas with highest tumor grade. Regular slides were used for the skeletal metastases.

