## Supplementary Figures and Tables for "PRSS2 stimulates tumor growth by remodeling the TME via repression of Tsp1"

Figure S1.

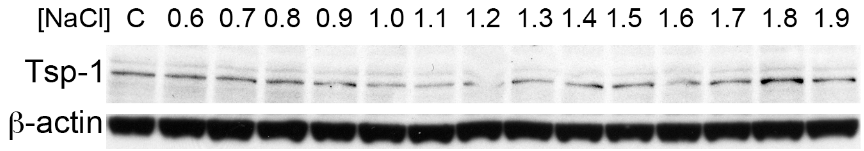


Western blot of WI38 fibroblasts treated with fractions of PC3M-LN4 conditioned media eluted from a heparin-sepharose+$Cu^{2+}$ column with a linear gradient of NaCl plus 20mM imidazole.

**Supplementary Figure S2.**

List of proteins present in Tsp-1 repressing fractions (1.0 and 1.1M) and inactive adjacent fractions

**0.9M NaCl**

- ~~Keratin 9~~
- ~~Keratin 1~~
- Keratin, CK2
- ~~Keratin, CK 10~~
- Keratin 10
- Keratin, CK6a
- Keratin, CK14
- Keratin, CK6e
- Keratin, CK5
- Keratin, CK16
- Cytokeratin type II
- Hornerin
- GAPDH
- Histone H2A.m
- Keratin, CK15
- 49 kDa protein
- Histone H2B.q
- ALB protein
- Keratin K6irs
- Lactotransferrin ~~precursor~~
- Histone H4
- ~~Serotransferrin precursor~~
- keratin 25 irs1
- Hypothetical protein DKFZp686J1375
- Hypothetical protein DKFZp686B2031
- ZF 263
- ~~MAP3K12~~
- Ciliary rootlet coiled-coil, rootletin
- XA protein
- Splice Isoform HMW of Kininogen-1 precursor
- Keratin, CK18
- Keratin, Ha6
- Cadherin protein
- Tigger transposable element derived 7
- Hypothetical protein FLJ90556

**1.0M NaCl**

- ~~Keratin, CK1~~
- ~~Keratin 9~~
- ~~Keratin, CK10~~
- ~~Keratin 2a~~
- ~~Lactotransferrin precursor~~
- ~~Keratin 1B~~
- ~~Serotransferrin precursor~~
- ~~ALB protein~~
- ~~similar to KIAA1501~~
- ~~24-dehydrocholesterol reductase precursor~~
- ~~Tropomodulin 1~~
- **Protease serine 2 isoform B**
- ~~Hypothetical protein FLJ90556~~
- ~~Splice Isoform 2 of WD-repeat protein 22~~
- ~~Ciliary rootlet coiled-coil, rootletin~~
- ~~EVH1 domain binding protein~~
- ~~MAP3K12~~
- ~~Hypothetical protein DKFZp434H152~~

**1.1M NaCl**

- ~~Keratin 9~~
- ~~Keratin 1~~
- ~~Keratin, CK10~~
- ~~Keratin, CK 2~~
- ~~Keratin, CK16~~
- ~~Keratin, CK6C~~
- ~~Keratin, CK14~~
- ~~Keratin, CK5~~
- ~~GAPDH~~
- ~~Keratin, CK13~~
- ~~Histone H2A.m~~
- ~~Histone H2B.q~~
- ~~Lactotransferrin precursor~~
- ~~Keratin, Hb4~~
- ~~Serotransferrin precursor~~
- ~~Histone 1, H2aa~~
- ~~Hypothetical protein LOC65250~~
- ~~DKFZp686J1375~~
- ~~Desmoglein-1~~
- ~~Cadherin protein~~
- ~~similar to KIAA1501~~
- ~~MAP3K 12~~
- ~~Hypothetical protein DKFZp686B2031~~
- ~~ALB protein~~
- **Protease serine 2 isoform B**
- ~~Junction plakoglobin~~
- ~~similar to TP-beta~~
- ~~similar to CK18~~
- ~~PRO0650~~
- ~~Nuclear pore membrane glycoprotein 210-like~~

**1.2M NaCl**

- ~~Keratin 1~~
- ~~Keratin 9~~
- ~~Keratin, CK10~~
- 57 kDa protein
- Keratin 10
- Keratin, CK2
- Keratin 1B
- Keratin 6L
- Keratin, type I cytoskeletal 14
- Keratin 5c
- Keratin, CK3
- GAPDH
- ALB protein
- Hypothetical protein FLJ20261
- similar to KIAA1501
- similar to KRT8
- keratin 25 irs1
- ROK1
- Cadherin protein
- 26 kDa protein
- hypothetical protein ABC A13
- Tropomodulin 1

**Supplementary Figure S3.**

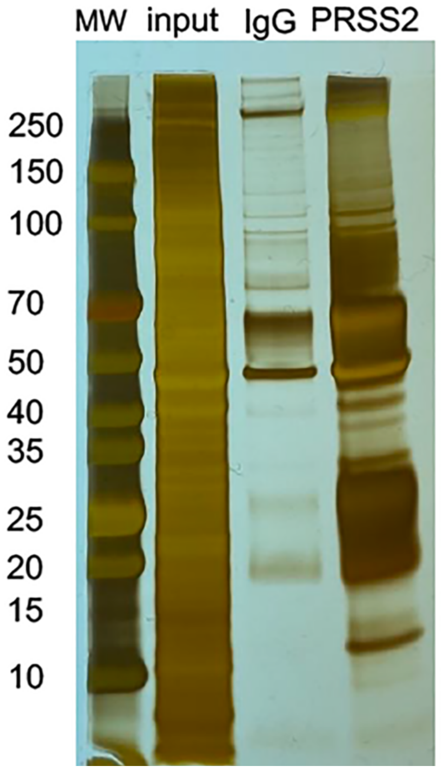


Silver stained polyacrylamide gel loaded with (from left to right) molecular weight markers (MW), input from IP experiment, immunoprecipitated from IgG control (IgG), and immunoprecipitated from $\alpha$-PRSS2 antibody (PRSS2).

**Supplementary Figure S4**
Liist of proteins immunoprecipitated only by PRSS2 that have more than 3 spectral counts

| | Spectral Counts | | Area Under the Curve |
|---|---|---|---|
| Gene Symbol | PRSS2 | IgG | |
| TRIM21 | 20 | 0 | 1890180000 |
| SVIL | 95 | 0 | 1596750000 |
| FLNC | 70 | 0 | 1306770000 |
| STRN3 | 38 | 0 | 1173520000 |
| PLEC | 112 | 0 | 892116000 |
| MYO18A | 26 | 0 | 612353000 |
| CTTNBP2NL | 39 | 0 | 597505000 |
| SPTAN1 | 64 | 0 | 524906000 |
| CORO1C | 9 | 0 | 478883000 |
| MPRIP | 42 | 0 | 461029000 |
| TJP1 | 49 | 0 | 431766000 |
| IGKV1D-13 | 5 | 0 | 380172000 |
| STOM | 16 | 0 | 377938000 |
| IQGAP1 | 49 | 0 | 366736000 |
| FLII | 25 | 0 | 361198000 |
| TPM1 | 12 | 0 | 332550000 |
| MACF1 | 117 | 0 | 331966000 |
| FLNB | 44 | 0 | 319585000 |
| CAPZB | 12 | 0 | 305767000 |
| PDIA6 | 7 | 0 | 302051000 |
| HLA-E | 5 | 0 | 277995000 |
| UBR5 | 51 | 0 | 248712000 |
| TJP2 | 33 | 0 | 238833000 |
| RAI14 | 36 | 0 | 236049000 |
| ACTN4 | 33 | 0 | 234127000 |
| CLTC | 23 | 0 | 232240000 |
| DKFZp686J1372 | 9 | 0 | 227785000 |
| LUZP1 | 39 | 0 | 219160000 |
| MYO6 | 25 | 0 | 209570000 |
| RRBP1 | 19 | 0 | 182661000 |
| NES | 31 | 0 | 179358000 |
| PPP1R18 | 19 | 0 | 172414000 |
| MYO5A | 29 | 0 | 171120000 |
| MYO1E | 17 | 0 | 170366000 |
| TMOD3 | 14 | 0 | 163159000 |
| TPM2 | 4 | 0 | 162652000 |
| CTTN | 18 | 0 | 153889000 |
| LGALS1 | 5 | 0 | 148607000 |
| SPECC1L | 31 | 0 | 137313000 |
| VDAC2 | 10 | 0 | 132468000 |
| ARPC2 | 11 | 0 | 125070000 |
| MVP | 16 | 0 | 124451000 |
| A2M | 5 | 0 | 112823000 |
| STRN4 | 13 | 0 | 111163000 |
| SQOR | 11 | 0 | 104457000 |
| RPL23 | 6 | 0 | 103869000 |
| ARPC1B | 6 | 0 | 99707900 |
| HNRNPM | 21 | 0 | 95921400 |
| C4A | 6 | 0 | 95083900 |
| ACTR2 | 16 | 0 | 92455600 |
| CAV1 | 3 | 0 | 92114000 |

| | | | |
|---|---|---|---|
| TPM1 | 5 | 0 | 87281700 |
| DOCK7 | 20 | 0 | 82621200 |
| RPL24 | 6 | 0 | 82406600 |
| LRRFIP2 | 13 | 0 | 77413500 |
| CAPZA2 | 12 | 0 | 63921100 |
| ACTR3 | 9 | 0 | 59103300 |
| ISG15 | 3 | 0 | 57779300 |
| WDR1 | 6 | 0 | 56617800 |
| CALM2 | 5 | 0 | 56010800 |
| SQSTM1 | 6 | 0 | 52663400 |
| DYNC1H1 | 34 | 0 | 50494600 |
| CEP170 | 19 | 0 | 50289800 |
| CAVIN1 | 16 | 0 | 49687100 |
| MYO10 | 20 | 0 | 49180700 |
| SLC25A3 | 4 | 0 | 46887300 |
| VDAC1 | 7 | 0 | 41187500 |
| ASPH | 9 | 0 | 40456000 |
| ITGB1 | 8 | 0 | 38116400 |
| ABLIM3 | 9 | 0 | 37103600 |
| SLC25A5 | 5 | 0 | 36571500 |
| AFAP1 | 10 | 0 | 36557600 |
| TMOD2 | 5 | 0 | 35883600 |
| P4HA1 | 7 | 0 | 34660500 |
| SPECC1 | 13 | 0 | 32873900 |
| RPS8 | 5 | 0 | 30367900 |
| RPL4 | 3 | 0 | 30306300 |
| SSFA2 | 22 | 0 | 30296100 |
| ARF4 | 3 | 0 | 30046800 |
| HLA-C | 6 | 0 | 29705000 |
| TRIOBP | 6 | 0 | 28547500 |
| RPS4X | 5 | 0 | 27045400 |
| TMPO | 19 | 0 | 26906000 |
| PPP1R9B | 10 | 0 | 26658200 |
| VDAC3 | 7 | 0 | 25915800 |
| TGM2 | 4 | 0 | 24920400 |
| STRIP1 | 7 | 0 | 24646100 |
| FN1 | 15 | 0 | 24492200 |
| TUFM | 8 | 0 | 24353500 |
| SLMAP | 13 | 0 | 23954900 |
| LASP1 | 6 | 0 | 23527700 |
| DDX3X | 8 | 0 | 23481400 |
| COLGALT1 | 7 | 0 | 21826400 |
| RPS3 | 6 | 0 | 21716700 |
| DLST | 7 | 0 | 20524300 |
| RAB3B | 4 | 0 | 20460600 |
| PPP1R12C | 5 | 0 | 20170900 |
| COL6A3 | 18 | 0 | 19681200 |
| TFG | 9 | 0 | 19591300 |
| RPN2 | 5 | 0 | 18811600 |
| DNAJA1 | 5 | 0 | 18778200 |
| COPA | 11 | 0 | 18611900 |
| EEF2 | 5 | 0 | 18530900 |
| PABPC1 | 4 | 0 | 17596400 |
| EIF4A1 | 5 | 0 | 17024200 |
| PFKP | 7 | 0 | 16954700 |
| PPP1CA | 5 | 0 | 16400800 |

| | | | |
|---|---|---|---|
| KIF5B | 11 | 0 | 16181400 |
| RPS6 | 3 | 0 | 14874800 |
| RUVBL2 | 6 | 0 | 14861700 |
| CEMIP | 4 | 0 | 14598400 |
| SSH1 | 7 | 0 | 14351300 |
| EMD | 3 | 0 | 14309300 |
| GJA1 | 7 | 0 | 14288400 |
| DST | 26 | 0 | 14015100 |
| LGALS3BP | 7 | 0 | 13858700 |
| DNAJB11 | 4 | 0 | 13662600 |
| ABCD3 | 4 | 0 | 13336700 |
| LRRFIP1 | 6 | 0 | 13291900 |
| RPL7 | 4 | 0 | 13260900 |
| NEK9 | 4 | 0 | 13013300 |
| DNAJA2 | 3 | 0 | 12914200 |
| RAB11FIP1 | 4 | 0 | 12061600 |
| IGLC3 | 3 | 0 | 11999800 |
| FLOT2 | 7 | 0 | 11922500 |
| ACOT9 | 6 | 0 | 11745800 |
| DDX17 | 5 | 0 | 11584100 |
| RNF213 | 11 | 0 | 11515000 |
| SH3BP4 | 6 | 0 | 11479900 |
| TMOD1 | 8 | 0 | 11261700 |
| JCAD | 11 | 0 | 11058500 |
| RPS3A | 3 | 0 | 11047700 |
| ERLIN1 | 4 | 0 | 11025000 |
| EPB41L2 | 9 | 0 | 10941400 |
| STRN | 9 | 0 | 10619400 |
| CLIC1 | 4 | 0 | 10613800 |
| ERLIN2 | 4 | 0 | 10555900 |
| GANAB | 10 | 0 | 10505600 |
| PLOD1 | 4 | 0 | 10267600 |
| MAP1B | 15 | 0 | 10151200 |
| IGF2BP3 | 9 | 0 | 10147100 |
| RACGAP1 | 5 | 0 | 10029100 |
| XRCC5 | 4 | 0 | 9820490 |
| MYOF | 10 | 0 | 9712970 |
| RAB13 | 3 | 0 | 9618030 |
| CYB5R3 | 3 | 0 | 9409190 |
| RPL18 | 5 | 0 | 9387850 |
| ITPR3 | 11 | 0 | 9235000 |
| RPS18 | 3 | 0 | 9210520 |
| RRAS2 | 3 | 0 | 9199800 |
| ATP1A1 | 9 | 0 | 9158490 |
| PCBP1 | 4 | 0 | 8973260 |
| KANK2 | 5 | 0 | 8918050 |
| EPRS | 7 | 0 | 8795920 |
| DARS | 6 | 0 | 8677180 |
| CKAP4 | 6 | 0 | 8643520 |
| PCBP2 | 7 | 0 | 8564960 |
| RPL13 | 3 | 0 | 8531740 |
| AASS | 3 | 0 | 8401660 |
| DDOST | 4 | 0 | 8388800 |
| F5 | 3 | 0 | 8369430 |
| RUVBL1 | 3 | 0 | 8290570 |
| ATP2B1 | 6 | 0 | 8234260 |

| | | | |
|---|---|---|---|
| FLOT1 | 7 | 0 | 8156840 |
| PLOD3 | 4 | 0 | 8048030 |
| B2M | 3 | 0 | 7942520 |
| DLAT | 4 | 0 | 7906180 |
| CAD | 7 | 0 | 7813460 |
| SRGAP2 | 3 | 0 | 7702120 |
| STAT1 | 6 | 0 | 7320940 |
| PABPC4 | 6 | 0 | 7286940 |
| FAM120A | 6 | 0 | 7232780 |
| GTF2I | 7 | 0 | 7217940 |
| MICAL2 | 5 | 0 | 7013130 |
| LIMCH1 | 7 | 0 | 6872990 |
| ATP6V1A | 7 | 0 | 6803790 |
| DDX20 | 5 | 0 | 6592370 |
| LMOD1 | 4 | 0 | 6303040 |
| MX1 | 4 | 0 | 6204520 |
| RAB7A | 5 | 0 | 6166240 |
| EHD2 | 3 | 0 | 6151180 |
| THY1 | 3 | 0 | 6104270 |
| RPL5 | 5 | 0 | 5962340 |
| ATP2A2 | 5 | 0 | 5955970 |
| IARS | 5 | 0 | 5901710 |
| KLC1 | 6 | 0 | 5843160 |
| KIF23 | 7 | 0 | 5812760 |
| HADHB | 3 | 0 | 5811820 |
| COPB2 | 7 | 0 | 5771450 |
| FST | 4 | 0 | 5657020 |
| TGFBI | 4 | 0 | 5598210 |
| AHNAK2 | 5 | 0 | 5375960 |
| AP2B1 | 3 | 0 | 5358950 |
| RPL10 | 3 | 0 | 5332270 |
| NSF | 6 | 0 | 5324620 |
| ZC3HAV1 | 5 | 0 | 5284610 |
| CAMK2D | 5 | 0 | 5158590 |
| TLN1 | 8 | 0 | 5155710 |
| TCIRG1 | 3 | 0 | 5112590 |
| ICAM1 | 3 | 0 | 5099330 |
| MICAL3 | 6 | 0 | 5067530 |
| ADD1 | 6 | 0 | 5031700 |
| MAP7D1 | 7 | 0 | 4925080 |
| CSRP1 | 3 | 0 | 4903760 |
| RPLP0 | 3 | 0 | 4852000 |
| XRCC6 | 3 | 0 | 4817710 |
| PRKDC | 6 | 0 | 4768880 |
| PDIA4 | 4 | 0 | 4748170 |
| DNAJC13 | 10 | 0 | 4619970 |
| IFIT1 | 6 | 0 | 4453780 |
| DYNC1I2 | 3 | 0 | 4380370 |
| CYR61 | 3 | 0 | 4375140 |
| PSMD3 | 6 | 0 | 4325630 |
| TRPV2 | 3 | 0 | 4316930 |
| ARHGEF17 | 7 | 0 | 4242570 |
| PALLD | 5 | 0 | 4154950 |
| MSN | 3 | 0 | 4100780 |
| PDIA3 | 5 | 0 | 4065620 |
| DCTN4 | 4 | 0 | 4051850 |

| | | | |
|---|---|---|---|
| PSMC2 | 4 | 0 | 3867790 |
| SIPA1 | 4 | 0 | 3863240 |
| PLOD2 | 3 | 0 | 3771340 |
| ARF3 | 3 | 0 | 3752890 |
| PPP2R1A | 3 | 0 | 3720030 |
| DLG5 | 4 | 0 | 3713710 |
| LRCH1 | 3 | 0 | 3596760 |
| ITGA2 | 3 | 0 | 3584370 |
| ADD3 | 5 | 0 | 3507660 |
| XIRP1 | 5 | 0 | 3460690 |
| FILIP1L | 5 | 0 | 3452440 |
| HSD17B4 | 5 | 0 | 3331970 |
| MYO9B | 5 | 0 | 3259110 |
| MATR3 | 8 | 0 | 3254420 |
| ARHGDIA | 3 | 0 | 3183600 |
| IMMT | 3 | 0 | 3180020 |
| RASAL2 | 8 | 0 | 3168500 |
| AIFM1 | 4 | 0 | 3166970 |
| SYNPO | 5 | 0 | 3133090 |
| ARHGAP21 | 5 | 0 | 3065580 |
| ATAD3A | 4 | 0 | 3063610 |
| P4HA2 | 3 | 0 | 3042140 |
| PTX3 | 5 | 0 | 3006770 |
| SHCBP1 | 3 | 0 | 2937870 |
| HADHA | 5 | 0 | 2888940 |
| CTNND1 | 3 | 0 | 2801550 |
| CACNA2D1 | 7 | 0 | 2800170 |
| RFTN1 | 4 | 0 | 2769760 |
| AKAP8L | 4 | 0 | 2721010 |
| ERP44 | 4 | 0 | 2695210 |
| CTSB | 3 | 0 | 2660010 |
| GFPT1 | 6 | 0 | 2632190 |
| PAWR | 4 | 0 | 2599850 |
| UGGT1 | 4 | 0 | 2568850 |
| RNH1 | 3 | 0 | 2566020 |
| SSR1 | 3 | 0 | 2555350 |
| EDIL3 | 4 | 0 | 2518560 |
| TOM1 | 4 | 0 | 2484870 |
| AP2A1 | 3 | 0 | 2473910 |
| EHD1 | 3 | 0 | 2447210 |
| PEAK1 | 6 | 0 | 2423890 |
| COPB1 | 4 | 0 | 2391190 |
| ATP6V0A1 | 3 | 0 | 2388180 |
| MLEC | 6 | 0 | 2386570 |
| STRAP | 5 | 0 | 2314460 |
| SMTN | 5 | 0 | 2266670 |
| MX2 | 3 | 0 | 2261420 |
| OGDH | 3 | 0 | 2212360 |
| U2AF2 | 3 | 0 | 2139340 |
| CANX | 3 | 0 | 2092510 |
| IGKC | 3 | 0 | 2087520 |
| ACSL3 | 3 | 0 | 2057170 |
| RAB11FIP5 | 4 | 0 | 2024340 |
| STT3A | 3 | 0 | 2016430 |
| STRIP2 | 3 | 0 | 1992390 |
| TMEM43 | 3 | 0 | 1946820 |

| | | | |
|---|---|---|---|
| RHOG | 3 | 0 | 1852410 |
| PDLIM7 | 3 | 0 | 1843050 |
| EHD4 | 3 | 0 | 1834590 |
| UTRN | 6 | 0 | 1767060 |
| ARHGEF11 | 4 | 0 | 1745830 |
| PRRC2C | 3 | 0 | 1735810 |
| INF2 | 4 | 0 | 1726210 |
| PPFIBP1 | 3 | 0 | 1726090 |
| CALU | 3 | 0 | 1658870 |
| RAB34 | 3 | 0 | 1658440 |
| CAPN2 | 5 | 0 | 1644620 |
| TOM1L2 | 4 | 0 | 1616700 |
| PRRC2A | 4 | 0 | 1512690 |
| CPT1A | 3 | 0 | 1511920 |
| WARS | 4 | 0 | 1501560 |
| MAP4K4 | 3 | 0 | 1478240 |
| SLC38A2 | 4 | 0 | 1458710 |
| CCT6A | 3 | 0 | 1410580 |
| LRP1 | 3 | 0 | 1365810 |
| RARS | 3 | 0 | 1359700 |
| OSBPL3 | 6 | 0 | 1343330 |
| SLC25A1 | 4 | 0 | 1336370 |
| MAP4 | 3 | 0 | 1314150 |
| DKK3 | 5 | 0 | 1269030 |
| IGF2BP2 | 6 | 0 | 1245730 |
| CSDE1 | 3 | 0 | 1217850 |
| MAP1A | 3 | 0 | 1206720 |
| THBS1 | 3 | 0 | 1187540 |
| GAS2L1 | 4 | 0 | 1119960 |
| ENG | 3 | 0 | 1056210 |
| UGGT2 | 3 | 0 | 1049850 |
| SND1 | 3 | 0 | 1006040 |
| HNRNPL | 4 | 0 | 927958 |
| MYCBP2 | 5 | 0 | 912300 |
| IFIT3 | 4 | 0 | 838077 |
| DHX9 | 3 | 0 | 812430 |
| ILF3 | 3 | 0 | 753033 |
| SYNPO2 | 3 | 0 | 741934 |
| ETFA | 3 | 0 | 704462 |
| LAMC1 | 4 | 0 | 660796 |
| SYNCRIP | 3 | 0 | 652562 |
| GLS | 3 | 0 | 629496 |
| FASN | 4 | 0 | 627206 |
| UBA1 | 3 | 0 | 604135 |
| OSBPL8 | 3 | 0 | 456749 |
| ARL1 | 3 | 0 | 407510 |
| APPL2 | 3 | 0 | 401947 |
| ABCC1 | 3 | 0 | 370126 |
| SPAG9 | 3 | 0 | 286035 |
| TKT | 3 | 0 | 227483 |

**Supplementary Figure S5.**

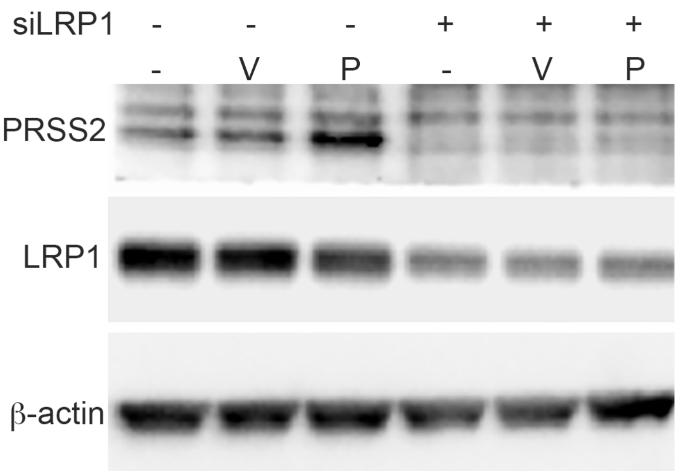


**PRSS2 is taken up by cells in an LRP1-dependent manner**
Western blot of PRSS2, LRP1, and actin in wild-type and siLRP1 transfected WI38 cells that were untreated (-) or treated with conditioned media from 293T cell transfected with empty pCMV-SPORT6 vector (V) or with pCMV-SPORT6-PRSS2 (P).

Supplementary Figure S6.

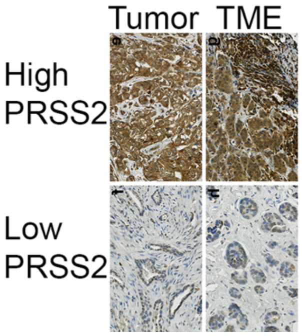


**PRSS2 expression in tumor and TME of breast cancer series 2.**

Supplementary Figure S7.

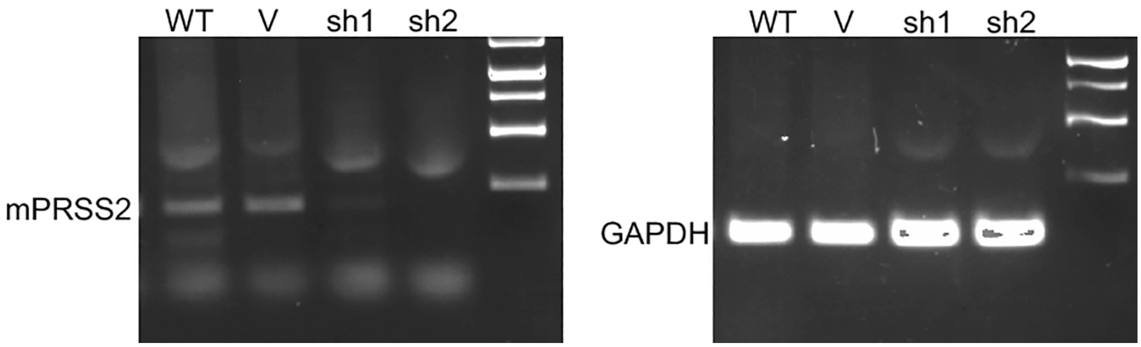


Ethydium bromide stained agarose gels loaded with (left) RT-PCR reactions for PRSS2 from WT, vector control (V), and two independent shPRSS2 transduced (sh1 and sh2) and (right) RT-PCR reactions for GAPDH from WT, vector control (V), and two independent shPRSS2 transduced (sh1 and sh2).

Supplementary Figure S8.

A.

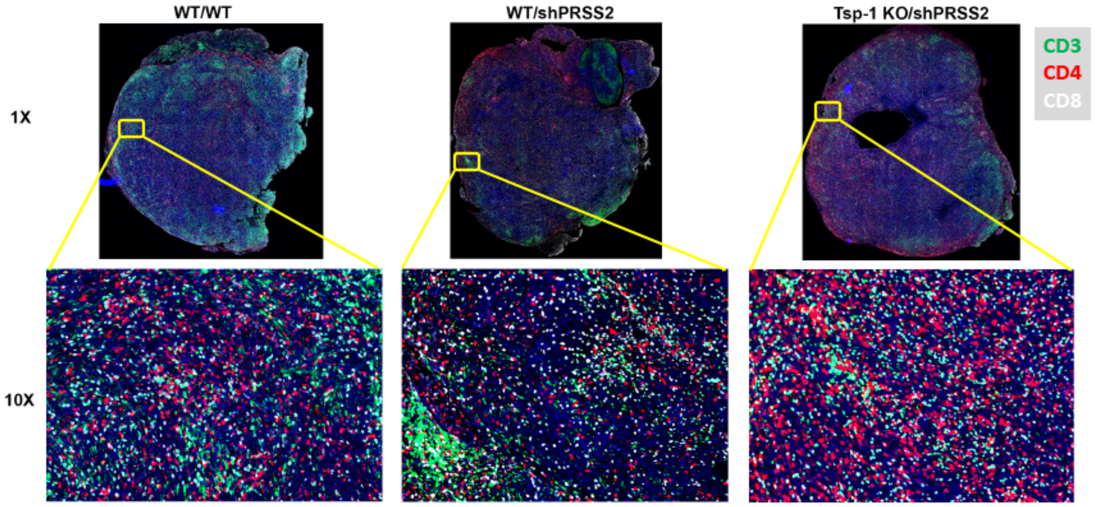


B.

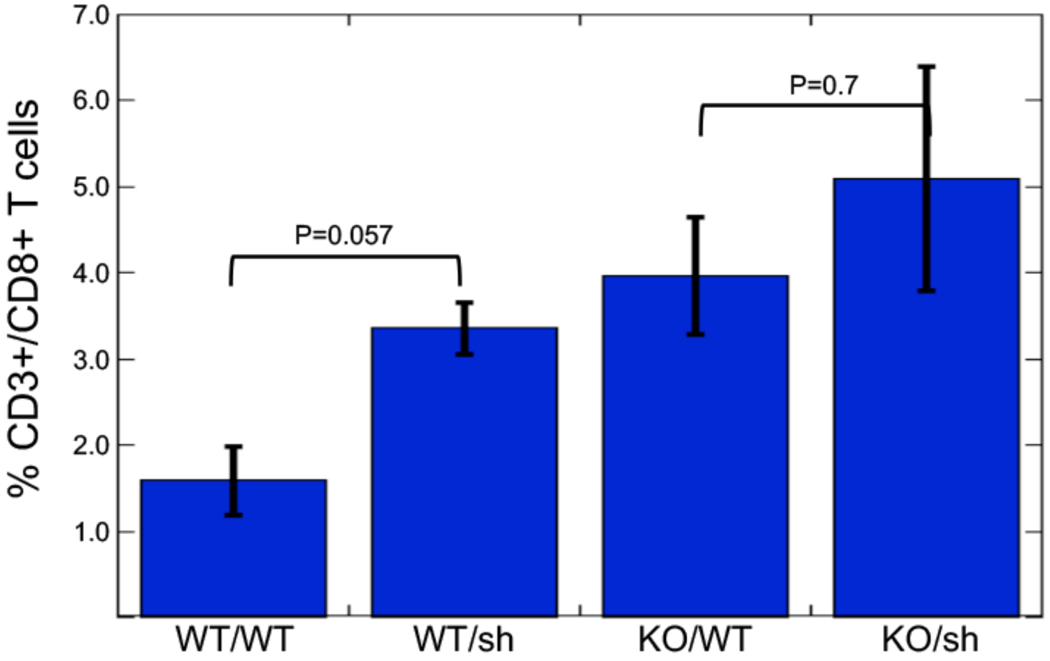


**A.** IHC staining of CD3, CD4 and CD8+ T cells in tumors formed by parental Pan02 tumor cells in wild-type C57Bl6/J mice (wt/wt), Pan02shPRSS2 cells in C57Bl6/J mice, and Pan02shPRSS2 cells in THBS1$^{-/-}$ C57Bl6/J mice.

**B.** Plot of average percentage of CD3+/CD8+ T cells in tumors formed by parental Pan02 tumor cells in wild-type C57Bl6/J mice (wt/wt), Pan02shPRSS2 cells in C57Bl6/J mice, and Pan02shPRSS2 cells in THBS1$^{-/-}$ C57Bl6/J mice.

Supplementary Figure S9.

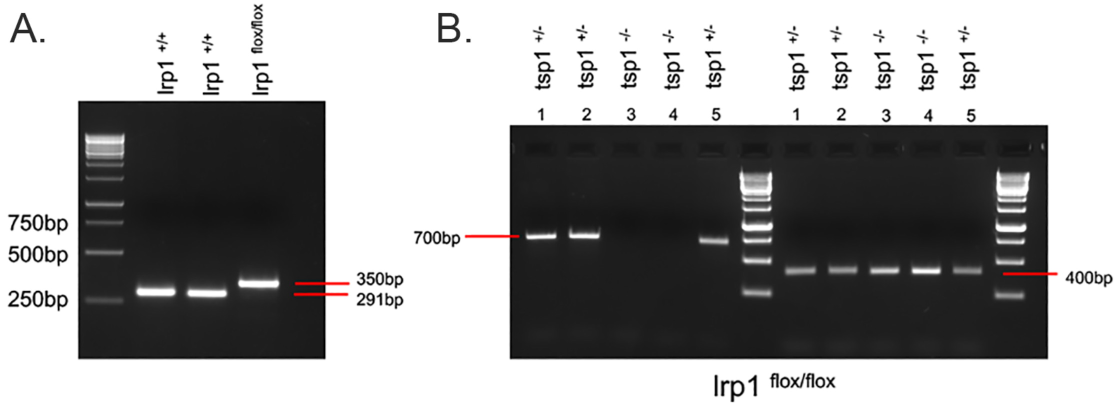


Ethydium bromide stained agarose gels of PCR products from genotyping experiments to confirm:

(A) Myeloid specific knockout of LRP1 as indicated by the presence of the 350bp product and absence of the 291bp product

(B) Knockout of Tsp-1 in 5 mice as indicated by the presence of the 400bp band and the absence of the 700bp band as observed in mice #3 and #4.

Supplementary Figure S10.

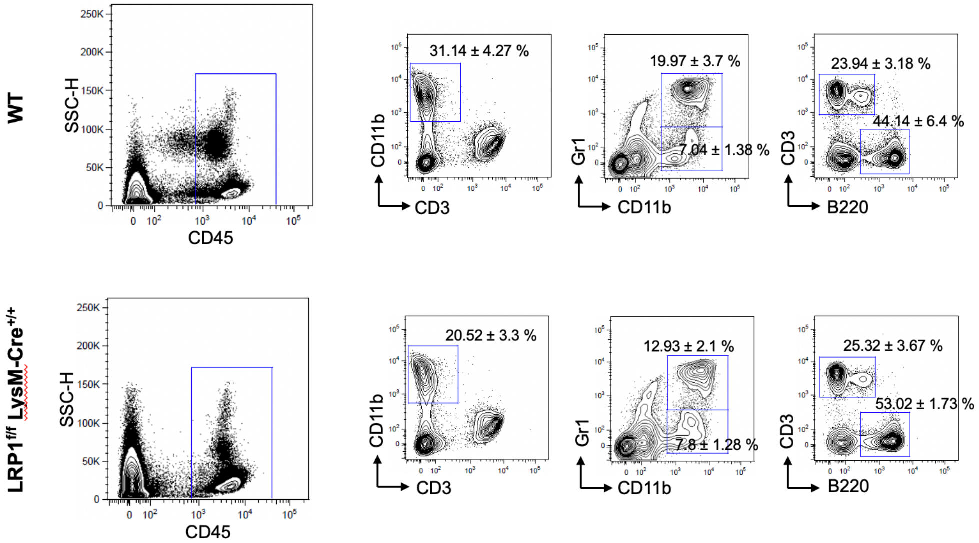


FACS analysis of myeloid and lymphoid cell in WT and LysM-Cre-LRP1$^{f/f}$ mice.

**Supplementary Table 1.** Associations between PRSS2 expression and selected features in breast cancer (series 2).

| Variables | PRSS2 EPITHELIUM | | | | | PRSS2 STROMA | | | | |
|---|---|---|---|---|---|---|---|---|---|---|
| | Low (n=114) | High (n=58) | OR | 95% CI | p-value[a] | Low (n=130) | High (n=42) | OR | 95% CI | p-value[a] |
| | n (%) | n (%) | | | | n (%) | n (%) | | | |
| **Histologic grade[b]** | | | | | <0.001 | | | | | <0.001 |
| Grade 1-2 | 66 (78.6) | 18 (21.4) | 1.0 | | | 74 (88.1) | 10 (11.9) | 1.0 | | |
| Grade 3 | 37 (48.1) | 40 (51.9) | 3.96 | 2.00, 7.88 | | 48 (62.3) | 29 (32.7) | 4.47 | 2.00, 10.00 | |
| **ER** | | | | | 0.516 | | | | | 0.006 |
| Pos (≥10%) | 61 (68.5) | 28 (31.5) | 1.0 | | | 75 (84.3) | 14(15.7) | 1.0 | | |
| Neg (<10%) | 53 (63.9) | 30 (36.1) | 1.23 | 0.66, 2.32 | | 55 (66.3) | 28 (33.7) | 2.73 | 1.32, 5.66 | |
| **Mitotic count[b]** | | | | | 0.001 | | | | | 0.004 |
| Low, ≤12.2 | 85 (71.4) | 34 (28.6) | 1.0 | | | 97 (81.5) | 22 (18.5) | 1.0 | | |
| High, >12.2 | 18 (42.9) | 24 (57.1) | 3.33 | 1.61, 6.91 | | 25 (59.5) | 17 (40.5) | 3.00 | 1.39, 6.48 | |
| **CK5/6[c]** | | | | | 0.439 | | | | | 0.305 |
| Neg, score=0 | 90 (68.2) | 42 (31.8) | 1.0 | | | 102 (77.3) | 30 (22.7) | 1.0 | | |
| Pos, score>0 | 24 (61.5) | 15 (38.5) | 1.34 | 0.64, 2.81 | | 27 (69.2) | 12 (30.8) | 1.51 | 0.68, 3.34 | |
| **p53** | | | | | 0.029 | | | | | 0.002 |
| Low, score ≤3 | 87 (71.3) | 35 (28.7) | 1.0 | | | 100 (82.0) | 22 (18.0) | 1.0 | | |
| High, score >3 | 27 (54.0) | 23 (46.0) | 2.12 | 1.07, 4.18 | | 30 (60.0) | 20 (40.0) | 3.03 | 1.46, 6.29 | |
| **pMVD[d]** | | | | | 0.721 | | | | | 0.178 |
| Low (< 1.45) | 86 (67.2) | 42 (32.8) | 1.0 | | | 99 (77.3) | 29 (22.7) | 1.0 | | |
| High (≥ 1.45) | 25 (64.1) | 14 (35.9) | 1.15 | 0.54, 2.43 | | 26 (66.7) | 13 (33.3) | 1.71 | 0.78, 3.74 | |
| ***BRCA1* mutation** | | | | | 0.124 | | | | | 0.359 |
| Absent | 89 (69.5) | 39 (30.5) | 1.0 | | | 99 (77.3) | 29 (22.7) | 1.0 | | |
| Present | 25 (56.8) | 19 (43.2) | 1.73 | 0.86, 3.51 | | 31 (70.5) | 13 (29.5) | 1.43 | 0.66, 3.09 | |

Series 2 (n=202). n: number of patients; OR: odds ratio; CI: confidence interval; ER: estrogen receptor; CK5/6: cytokeratin 5/6; pMVD: proliferative microvessel density.

[a] Pearson's chi-squared test.

[b] Eleven cases lack information on histologic grade and mitotic count (mitoses/mm2).

[c] One case lacks information on CK5/6 status.

[d] Five cases lack information on pMVD (FactorVIII+/Ki67+ vessels).
